# The mitotic protein NuMA plays a spindle-independent role in nuclear formation and mechanics

**DOI:** 10.1101/2020.05.02.070680

**Authors:** Andrea Serra-Marques, Ronja Houtekamer, Dorine Hintzen, John T. Canty, Ahmet Yildiz, Sophie Dumont

## Abstract

Eukaryotic cells typically form a single, round nucleus after mitosis, and failures to do so can compromise genomic integrity. How mammalian cells form such a nucleus remains incompletely understood. NuMA is a spindle protein whose disruption results in nuclear fragmentation. What role NuMA plays in nuclear integrity, or whether its perceived role stems from its spindle function, is unclear. Here, we use live imaging to demonstrate that NuMA plays a spindle-independent role in forming a single, round nucleus. NuMA keeps the decondensing chromosome mass compact at mitotic exit, and promotes a mechanically robust nucleus. NuMA’s C-terminus binds DNA in vitro and chromosomes in interphase, while its coiled-coil acts as a regulatory and structural hub: it prevents NuMA from binding chromosomes at mitosis, regulates its nuclear mobility and is essential for nuclear formation. Thus, NuMA plays a long-range structural role in building and maintaining an intact nucleus, as it does for the spindle, playing a protective role over the cell cycle.

## Introduction

Most eukaryotic cells enclose their genome within a single, round nucleus. When chromosomes are not properly incorporated into the nucleus, they can form micronuclei which can be vulnerable to DNA damage and chromothripsis, leading to genomic instability and disease (Andreassi et al., 2011; Crasta et al., 2012; Hatch et al., 2013; Zhang et al., 2015). Further, while different cell types have nuclei of different shapes, within a given cell type nuclear shape is regulated and stereotypical, and nuclear shape defects can correlate with altered gene expression, disease and aging (Le Berre et al., 2012; Dahl et al., 2008; Skinner and Johnson, 2017; Thomas et al., 2002). While some of the mechanisms required for nuclear formation are known, how mammalian cells pack many chromosomes into a single nucleus with a simple rounded shape is not well understood. More broadly, understanding how nm-scale molecules give rise to higher order μm-scale structures of a given number and shape remains a frontier.

Several mechanisms are known to promote the formation of a single nucleus and to give rise to its shape. For example, bringing anaphase chromosomes into a tight cluster at mitosis promotes the formation of a single nucleus. Indeed, erroneous chromosome-to-spindle attachments (Cimini et al., 2004; Liu et al., 2018; Liu and Pellman, 2020; Thompson and Compton, 2011), misregulated centrosome amplification (Ganem et al., 2009), and defective metaphase chromosome alignment and anaphase compaction (Fonseca et al., 2019) can all lead to micronucleation. At late mitosis, the protein BAF builds a dense chromatin layer around the anaphase chromosome mass, limiting membrane penetration to prevent the formation of multiple nuclei (Samwer et al., 2017). Similarly, diverse factors are thought to influence nuclear shape. These range from the shape of the anaphase chromosome mass (Fonseca et al., 2019; Mora-Bermúdez et al., 2007) and chromatin organization and compaction (Furusawa et al., 2015; Stephens et al., 2017) to the mechanics of the nuclear lamina, cytoskeleton and extracellular environment (Buxboim et al., 2010; Chu et al., 2017; Dahl et al., 2008; Lammerding et al., 2005).

Despite what we know about nuclear formation, understanding when – at mitosis versus at nuclear formation – and how directly certain molecules play a role in this process remains challenging. The protein <u>Nu</u>clear <u>M</u>itotic <u>A</u>pparatus (NuMA), for example, is essential to spindle assembly (Compton and Luo, 1995; Kallajoki et al., 1991; Yang and Snyder, 1992) and chromosome segregation (Silk et al., 2009; Haren et al., 2009), as well as to the formation of a single nucleus (Compton and Cleveland, 1993; Kallajoki et al., 1991, 1993). NuMA contains two globular domains, its N- and C-termini, connected through seven coiled-coil domains (Compton et al., 1992; Yang et al., 1992). At mitosis, NuMA localizes to the spindle and cell cortex where it drives spindle structural stability and orientation, respectively (Compton et al., 1992; Kallajoki et al., 1991; Kiyomitsu and Cheeseman, 2012; Kotak et al., 2013). At interphase, NuMA localizes to the nucleus. It has been proposed to have a post-mitotic role in nuclear formation (Compton and Cleveland, 1993; Kallajoki et al., 1991, 1993), which could be a direct or indirect role (Cleveland, 1995; Kallajoki et al., 1991, 1993; Merdes and Cleveland, 1998), and to promote DNA repair and chromatin organization in the nucleus (Abad et al., 2007; Kivinen et al., 2010; Vidi et al., 2014). Further, NuMA has been reported to self-assemble into a filamentous network in the nucleus (Gueth-Hallonet et al., 1998; Zeng et al., 1994), and to directly bind DNA *in vitro* (Ludérus et al., 1994). However, NuMA’s essential function at mitosis has made it difficult to determine whether it also acts at nuclear formation independently of its spindle function. If it does, we do not know what this nuclear formation function is, or how NuMA’s structure supports it and regulates it over the cell cycle.

Here, we show that NuMA is essential for the formation of a single, round nucleus in human cells independent of its spindle role. We find that in the absence of NuMA nuclear defects appear at the moment of nuclear envelope reassembly, even without prior spindle defects. Further, we show that in cells lacking NuMA the chromosome mass expands faster at mitotic exit, and that nuclei are mechanically compromised. We demonstrate that NuMA’s coiled-coil is essential to forming a single nucleus and to promoting the formation of stable nuclear NuMA assemblies. Further, we find that NuMA’s coiled-coil prevents the premature binding of NuMA’s C-terminus – which we show directly binds DNA *in vitro* – to chromosomes at mitosis. Thus, NuMA’s coiled-coil serves as a hub that controls both its localization and function over the cell cycle. Together, these findings establish a spindle-independent, *bona fide* role for NuMA in nuclear formation and mechanics, and suggest that NuMA’s long coiled-coil helps it organize chromosomes over microns at interphase. More broadly, they highlight NuMA’s key long-range structural role throughout the cell cycle, protecting both the spindle and the nucleus, two of the largest structures in cells.

## Results

### NuMA plays a spindle-independent role in the formation of a single, round nucleus

To investigate whether NuMA contributes to the formation of the mammalian nucleus independently of its spindle function, we used inducible CRISPR/Cas9 RPE1 cells to knockout (KO) NuMA (Hueschen et al., 2017), and looked at nuclear formation after a spindle-less mitosis. To do so, we treated cells with nocodazole to depolymerize spindle microtubules, and with reversine to bypass the spindle assembly checkpoint. With this approach, mitotic cells do not assemble a spindle, exit mitosis and form a single nucleus of unsegregated chromosomes (Samwer et al., 2017). We induced NuMA KO cells at t=0 with doxycycline (DOX) to deplete endogenous NuMA (Fig. S1A), treated them with nocodazole and reversine after 72 h, and fixed and imaged them 18-24 h after drug addition (termed “96 h” onward for simplicity; Fig. 1A). As expected, control (uninduced) drug-treated cells did not assemble a spindle (Fig. S1B), and yet still formed a single, round (termed “normal”) nucleus after mitosis (Fig. 1B-C; Samwer et al., 2017), as did Cas9-induced cells without a sgRNA for NuMA (Fig. S1C).

**Figure 1.**
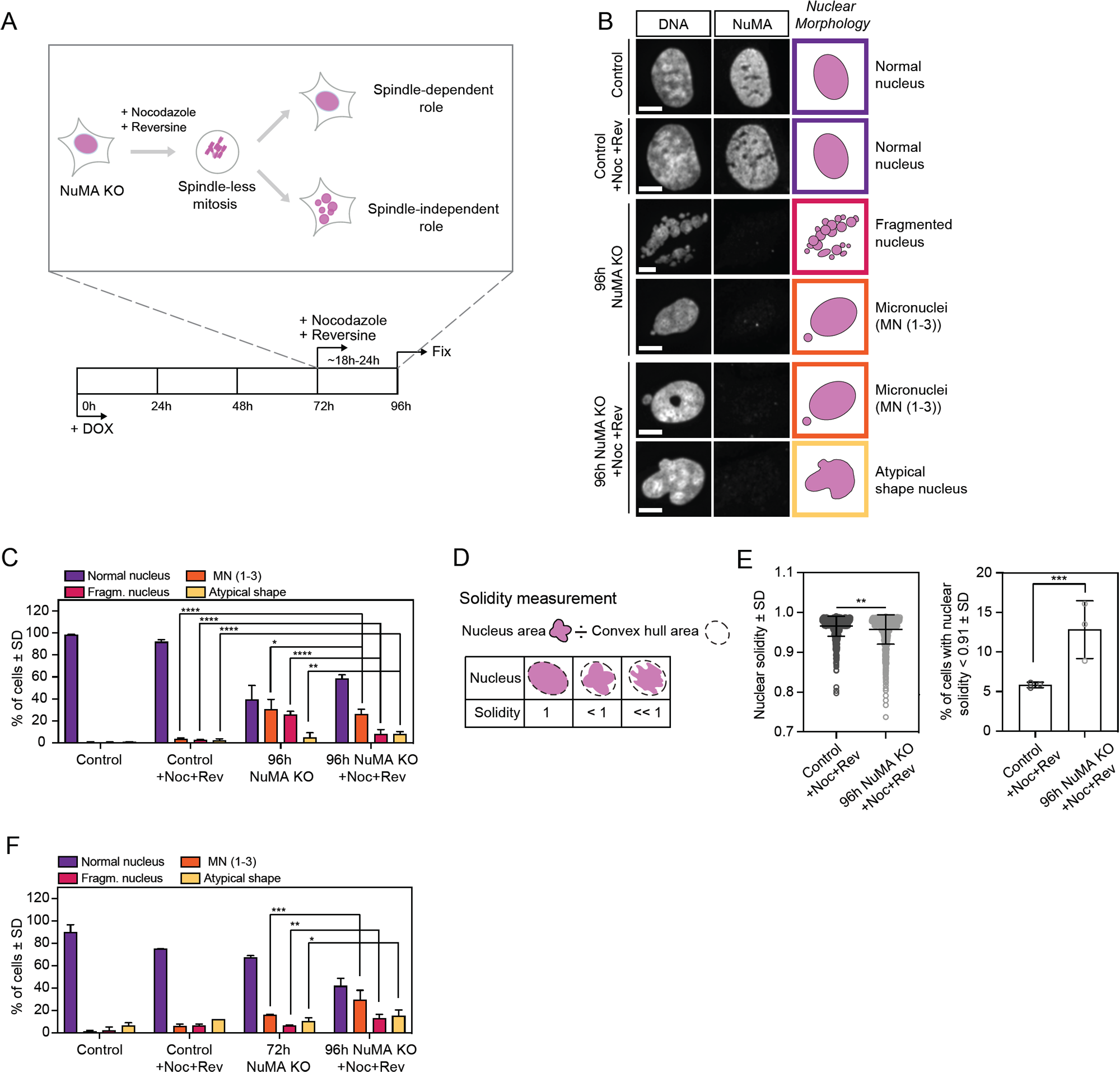
NuMA plays a spindle-independent role in the formation of a single, round nucleus. **(A)** Experimental design for inducible CRISPR/Cas9 NuMA KO RPE1 cells to undergo and exit a spindle-less mitosis. Endogenous NuMA is depleted by DOX addition starting at 0h, and cells treated with nocodazole (664 nM) and reversine (320 nM) for a spindle-less mitosis starting at 72h, and subsequently fixed and stained for immunofluorescence at 96h. **(B)** Representative immunofluorescence images of nuclear morphologies observed in uninduced NuMA KO RPE1 (control) cells, control cells treated with nocodazole and reversine (for 24h), 96h NuMA KO cells and 96h NuMA KO cells treated with nocodazole and reversine (for last 24h). Cells were stained for tubulin (not shown), DNA (Hoechst) and NuMA. Scale bar: 10 μm. **(C)** Percentage of cells with different nuclear morphologies in the indicated conditions, from DNA images in (B). n=1385 (control), 1123 (control +Noc+Rev), 638 (96h NuMA KO) and 649 (96h NuMA KO +Noc+Rev) cells, from 3 independent experiments. Fisher’s exact test: ****p<0.0001; **p=0.003; *p=0.01. The same nuclear morphologies are compared within different conditions. **(D)** The solidity of nuclei corresponds to the ratio of the nucleus’ area to its convex hull area. **(E)** Nuclear solidity in nocodazole and reversine treated control and 96h NuMA KO cells, and corresponding percentage of cells with nuclear solidity 2 standard deviations below the control mean. n= 750 (control) and 656 (96h NuMA KO) cells, from 3 independent experiments. Left plot, Mann-Whitney test: **p=0.001. Right plot, Fisher’s exact test: ***p=0.0001. **(F)** Percentage of cells with different nuclear morphologies in the indicated conditions, from DNA images. n=285 (control), 456 (control +Noc+Rev), 321 (72h NuMA KO) and 347 (96h NuMA KO +Noc+Rev) cells, from 2 independent experiments. Fisher’s exact test: ****p<0.0001; ***p=0.0001; **p=0.009; *p=0.03. The same nuclear morphologies are compared within different conditions.

In turn, NuMA KO cells with a defective spindle (untreated with drugs; Fig. S1B) presented nuclei with several fragments (Fig. 1B-C), as previously observed (Compton and Cleveland, 1993; Kallajoki et al., 1991, 1993). Additionally, these cells also presented micronuclei (1-3 small nuclei in addition to the primary nucleus) and atypically-shaped nuclei (Fig. 1B-C). In spindle-less NuMA KO cells (i.e. treated with nocodazole and reversine drugs), the fraction of cells with fragmented nuclei was significantly reduced, while the fraction of cells with micronuclei and atypically shaped nuclei remained similar (Fig. 1B-C). This suggests that fragmented nuclei result at least partly from spindle-related failures, and that micronucleation and atypically shaped nuclei result from spindle-independent failures. To quantitatively compare nuclear roundness, we measured the solidity of primary nuclei. Solidity significantly decreased in spindle-less NuMA KO cells compared to control cells (Fig. 1D-E), and decreased as NuMA depletion extent increased (Fig. S1D-E). Finally, because NuMA depletion occurred gradually over four days (Fig. S1A) we sought to determine whether nuclear defects appeared when spindles were present (before drug addition) or absent (after drug addition). To do so, we repeated the same assay (Fig. 1A) and compared cells fixed at 72 h, just before drug addition, and at 96 h, after drug treatment. The fraction of NuMA KO cells with fragmented nuclei, micronuclei, and atypically shaped nuclei all increased significantly in the 18-24 h spindle-less period (Fig. 1F). Together, these findings indicate that NuMA plays a *bona fide* role in the formation of a single and round nucleus, independent of its spindle function.

### NuMA acts at the time of nuclear formation, compacting the chromosome mass at mitotic exit

To gain insight into NuMA’s nuclear function, we sought to determine whether it is required before, after or at the time of nuclear formation. To do so, we performed live cell imaging of inducible NuMA KO RPE1 cells treated with nocodazole and reversine and stably expressing the histone marker mCherry-H2B and the nuclear envelope marker EGFP-Lap2β. As for fixed cells (Fig. 1), live imaging revealed a significant increase in the fraction of spindle-less NuMA KO cells with fragmented nuclei, micronuclei and atypically shaped nuclei at 96h compared to control cells (Fig. 2A-B). To define when NuMA is needed, we followed the trajectories of individual cells that entered mitosis with a single, round nucleus (‘mother cells’) until they exited spindle-less mitosis, formed a nucleus and beyond (‘daughter cells’) (Fig. 2C). From mother cells with normal nuclei, NuMA KO daughter cells exited mitosis with more fragmented nuclei, micronuclei and atypically shaped nuclei than control cells (Fig. 2D-E; Videos 1A-B). In these cell trajectories, we observed morphology defects appear as chromosomes decondensed and the nuclear envelope formed. We did not observe defects appear before nor soon after nuclear formation. To test whether NuMA could help keep the decondensing chromosome mass together (either directly or indirectly) as the nuclear envelope formed, we measured the expansion of this mass with and without NuMA. We found that the chromosome mass expanded faster, and further, at nuclear formation in NUMA KO cells than in control cells (Fig. 2F). Together, these findings indicate that NuMA’s spindle-independent role in building the nucleus takes place at the time of nuclear envelope formation, rather than before or after, and that NuMA keeps the chromosome mass compact during this time.

**Figure 2.**
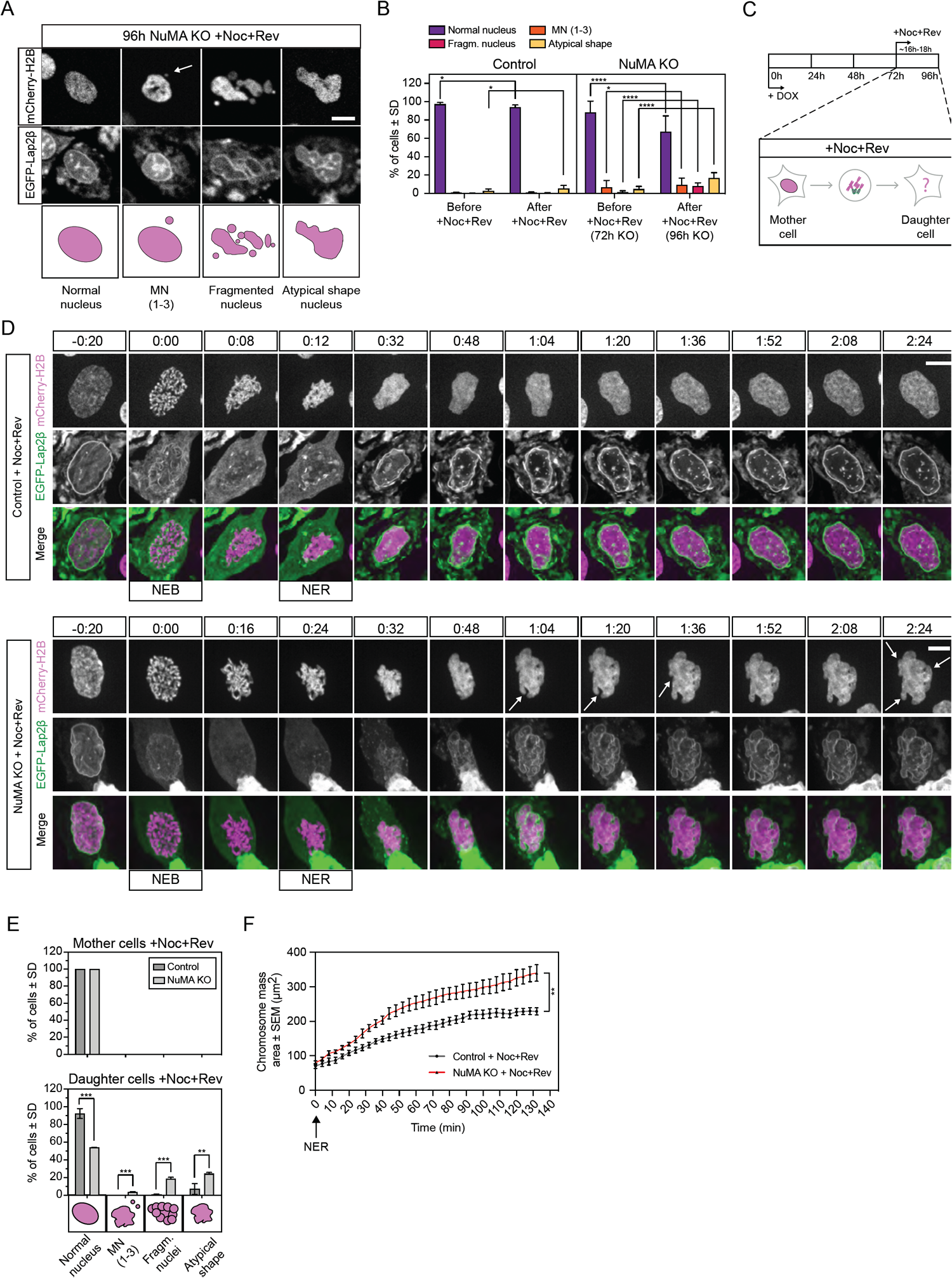
NuMA acts at the time of nuclear formation, compacting the chromosome mass at mitotic exit. **(A)** Representative live images of observed nuclear phenotypes after 16-18h of nocodazole and reversine treatment (spindle-less mitosis) in 96h NuMA KO RPE1 cells stably expressing mCherry-H2B and EGFP-Lap2β. Arrow indicates a micronucleus. Scale bar: 5 μm. **(B)** Percentage of cells with different nuclear morphologies before and 16-18h after nocodazole and reversine treatment (spindle-less mitosis). All cells, and not only those that went through mitosis, were analyzed. n=476 (control) and 820 (NuMA KO) cells from 5 and 2 independent experiments, respectively. Fisher’s exact tests: *p<0.01; ***p<0.0001. **(C)** Experimental design as in Fig. 1A, except that individual cell trajectories were followed by live imaging for the 16-18h period in nocodazole (664 nM) and reversine (control 1 μM; NuMA KO 320 nM). Mother cell, cell before mitotic entry; daughter cell, cell after mitotic exit. **(D)** Representative timelapse images of control and NuMA KO RPE1 cells stably expressing mCherry-H2B (magenta) and EGFP-Lap2β (green) that were followed live through spindle-less mitosis for 16-18h, noting nuclear envelope breakdown (NEB) and reformation (NER). White arrows indicate nuclear defects. Time in h:min. Scale bar: 10 μm. **(E)** Percentage of cells with the indicated nuclear phenotypes in uninduced control and NuMA KO cells treated with nocodazole and reversine just before mitotic entry (mother cells) into a spindle-less mitosis and just after mitotic exit (daughter cells). Mother cells include only cells that entered mitosis with a single, round nucleus. n=508 (control) and 308 (NuMA KO) cells from 5 and 2 independent experiments, respectively. One-way ANOVA: **p < 0.01; ****p < 0.0001). **(F)** Expansion of the chromosome mass from the time of nuclear envelope reformation (NER, t=0) in control and NuMA KO cells treated with nocodazole and reversine for a spindle-less mitosis. n=5 cells in each condition. Mann-Whitney test: **p=0.008 at 132 min.

### NuMA’s coiled-coil is required for the formation of a single nucleus and modulates its mobility in the nucleus

Electron microscopy and overexpression of NuMA coiled-coil truncations suggest that the coiled-coil helps form nuclear NuMA filaments, which could in principle contribute to nuclear stability (Gueth-Hallonet et al., 1998; Harborth et al., 1999; Zeng et al., 1994). To determine if NuMA’s coiled-coil is required for nuclear formation, we tested the ability of a truncation of NuMA that lacks most of the coiled-coil (NuMA-Bonsai) (Fig. 3A) to rescue the knockout of endogenous NuMA, when expressed at near endogenous levels (Fig. 3B-C; S2A). As a control, overexpression of similar levels of NuMA-FL-EGFP (full length, FL; Fig. S2A) in NuMA KO cells mostly rescued the nuclear defects observed in the absence of endogenous NuMA, even though some cells still presented micronuclei (Fig. 3B-C). In turn, NuMA-Bonsai-EGFP, which contains only a short portion of the coiled-coil, failed to significantly rescue micronucleation (Fig. 3B-C). This is consistent with NuMA-Bonsai being sufficient for NuMA’s role in the spindle body (Forth et al., 2014; Hueschen et al., 2017) but not at nuclear formation, with nuclear fragmentation coming largely from spindle defects, and micronucleation coming from spindle-independent defects, being revealed without spindle defects (Fig. 1C). Consistently, inducible NuMA KO cells stably expressing NuMA-Bonsai-EGFP (at higher levels than endogenous NuMA; Fig. S2B) had micronuclei (Fig. S2C) but no detectable chromosome segregation errors (Fig. S2D). Importantly, this indicates that NuMA’s nuclear formation role is not only essential after a drug-induced spindle-less mitosis, but also after normal mitosis. Together, these findings indicate that either all or part of NuMA’s coiled-coil is necessary for formation of a single nucleus.

**Figure 3:**
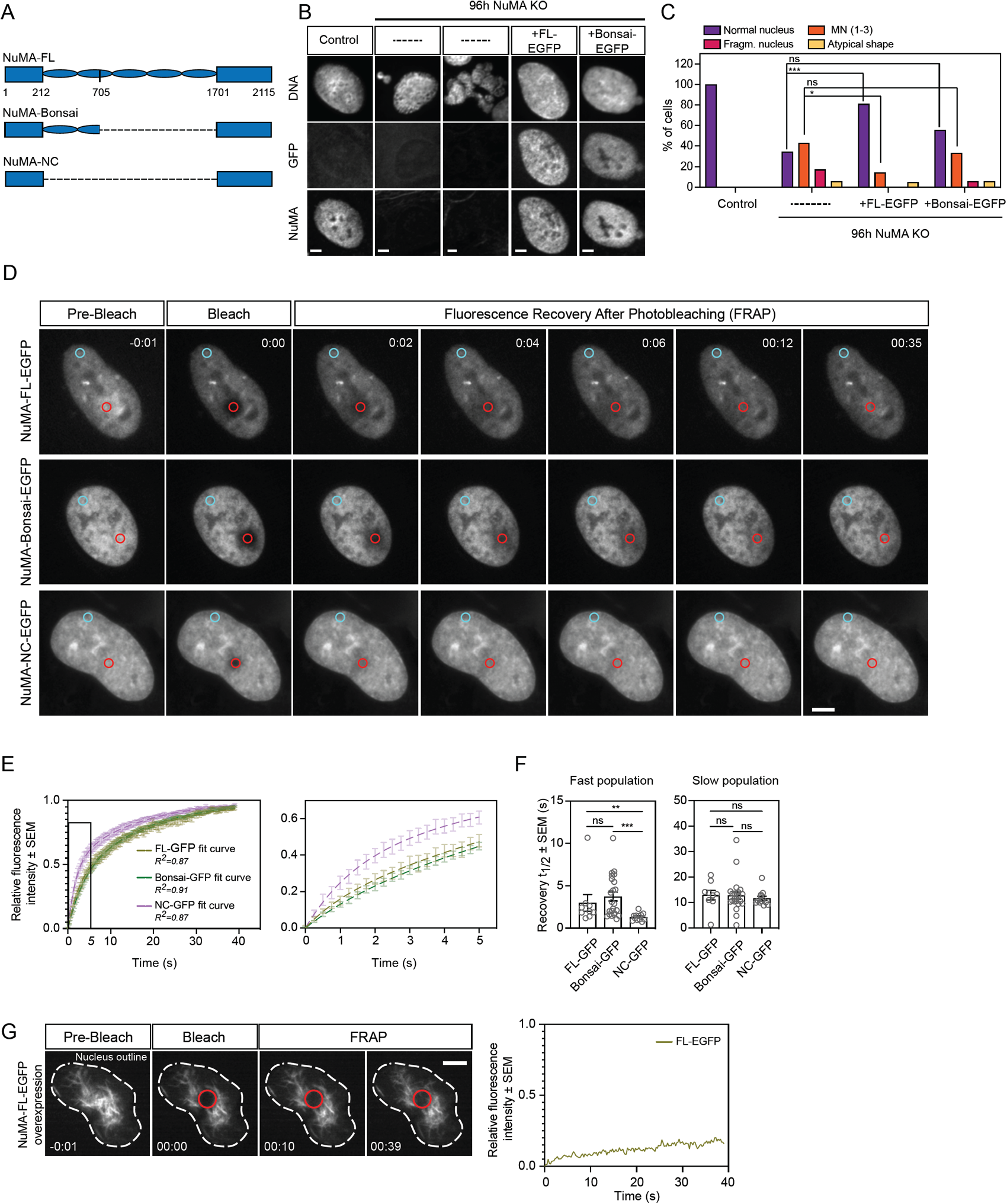
NuMA’s coiled-coil is required for the formation of a single nucleus and modulates its mobility in the nucleus. **(A)** Schematic representations of full-length (FL) NuMA and truncations NuMA-Bonsai and NuMA-NC, with amino acid numbers indicated. **(B)** Representative immunofluorescence images of nuclear morphologies observed in uninduced NuMA KO RPE1 cells (control), NuMA KO cells without exogenous constructs (--, two examples) and NuMA KO cells transiently expressing the rescue constructs NuMA-FL-EGFP or NuMA-Bonsai-EGFP. Cells were stained for tubulin (not shown), DNA (Hoechst), GFP and NuMA. Scale bar: 10 μm. **(C)** Percentage of cells with different nuclear morphologies observed in experiment from (B). n=38 (control), 35 (NuMA KO), 18 (NuMA KO + NuMA-FL-EGFP) and 21 (NuMA KO + NuMA-Bonsai-EGFP) cells. These rescues were repeated one more time, with similar results (not shown). Fisher’s exact test: ***p<0.0006; *p=0.04; ns, non-significant. **(D-E)** Fluorescence recovery after photobleaching (FRAP) of NuMA-FL-EGFP, NuMA-Bonsai-EGFP and NuMA-NC-EGFP in the nucleus of uninduced NuMA KO RPE1 cells. GFP intensity was measured in the bleached area (red circle) and in a non-bleached area (blue circle) to account for photobleaching. Scale bar: 10 μm. Time in min:s; 0:00 indicates the time of bleaching. n=11 (NuMA-FL-EGFP), 23 (NuMA-Bonsai-EGFP), and 12 (NuMA-NC-EGFP) cells. **(F)** Distribution of the fast and slow recovery halftimes (t_1/2_) of the different GFP-tagged NuMA proteins during FRAP. Mann Whitney test: **p=0.001; ***p<0.001 ns, non-significant. **(G)** Timelapse images showing the nucleus (dashed white line) of an uninduced NuMA KO RPE1 cell highly overexpressing NuMA-FL-EGFP and forming cable-like structures (left) and fluorescence recovery after photobleaching of the same cell (right). NuMA was bleached in the indicated red circle at 0:00 and only minimal GFP intensity recovered by 39 s. Time in min:s. Scale bar: 5 μm.

Given the role of NuMA’s coiled-coil in nuclear formation (Fig. 3B-C; S2C), and previous work (Gueth-Hallonet et al., 1998; Harborth et al., 1999; Zeng et al., 1994), we hypothesized that the coiled-coil helps form stable NuMA assemblies in the nucleus. To test this, we measured the turnover of NuMA-FL-GFP, NuMA-Bonsai-GFP, and NuMA-NC-GFP, a construct that lacks the entire coiled-coil, in the nucleus using fluorescence recovery after photobleaching (FRAP) (Fig. 3D-F). NuMA-FL-GFP and NuMA-Bonsai-GFP displayed similar recovery kinetics, while NuMA-NC-GFP recovered faster. Using a double exponential model, the fast half-life was shorter for NuMA-NC-GFP than NuMA-FL-GFP and NuMA-Bonsai-GFP, while the slow half-life was indistinguishable for all three constructs, independent of GFP-tagged protein expression levels within the analyzed range (Fig. S2E-F). Further, we observed that cells expressing the highest NuMA-FL-EGFP levels, excluded from the above analysis, can form stable cable-like nuclear structures that show much slower recovery after photobleaching (Fig. 3G; Video 2). This supports the notion that NuMA can self-assemble in the nucleus, at least in these conditions. In turn, we never observed such cable-like structures in NuMA-NC-EGFP cells, even at high expression levels. Together, these findings indicate that NuMA’s coiled-coil is necessary to nuclear formation, and suggest that it modulates NuMA’s mobility and may help it form a stable higher order structure in the nucleus.

### NuMA’s coiled-coil regulates chromatin binding by its C-terminus over the cell cycle

NuMA is only detected on chromosomes after initiation of nuclear envelope reassembly (Fig. S3), when it is imported into the nucleus (Compton et al., 1992). What mechanism ensures that NuMA only associates with chromosomes in interphase, when chromosomes are kept together, and not in mitosis when chromosomes must move apart? To begin answering this question, we asked how NuMA binds chromosomes in interphase, testing whether NuMA’s globular N- and C-terminal domains associate with chromosomes once in the nucleus. Both NuMA’s N- and C-termini were suggested to bind DNA (Radulescu and Cleveland, 2010), though no specific and direct comparisons were made. Using a light-controlled dissociation system, we sought to disconnect NuMA’s N- and C-termini from each other once they were both imported together into the nucleus. We engineered an opto-controllable NuMA-Bonsai protein (Opto-NuMA) using the LOVTRAP system (van Haren et al., 2018; Wang et al., 2016) (Fig. 4A). In the dark, when LOV2 and Zdk1 bind, NuMA-N(1-705)-PhusionRed-LOV2 and EGFP-Zdk1-NuMA-C localized to the nucleus, as expected since NuMA’s C-terminus contains a nuclear localization signal (NLS) (Tang et al., 1994). Upon blue light illumination, the nuclear localization of NuMA-C (imaged with the same wavelength that activates LOV2 and induces dissociation) remained unchanged. Meanwhile, NuMA-N became diffuse (Fig. 4B-C; Video 3), likely remaining in the nucleus due to its high molecular weight (123 kDa). Thus, whether chromatin binding is direct or indirect, NuMA’s C-terminus has a higher affinity for chromatin than its N-terminus.

**Figure 4.**
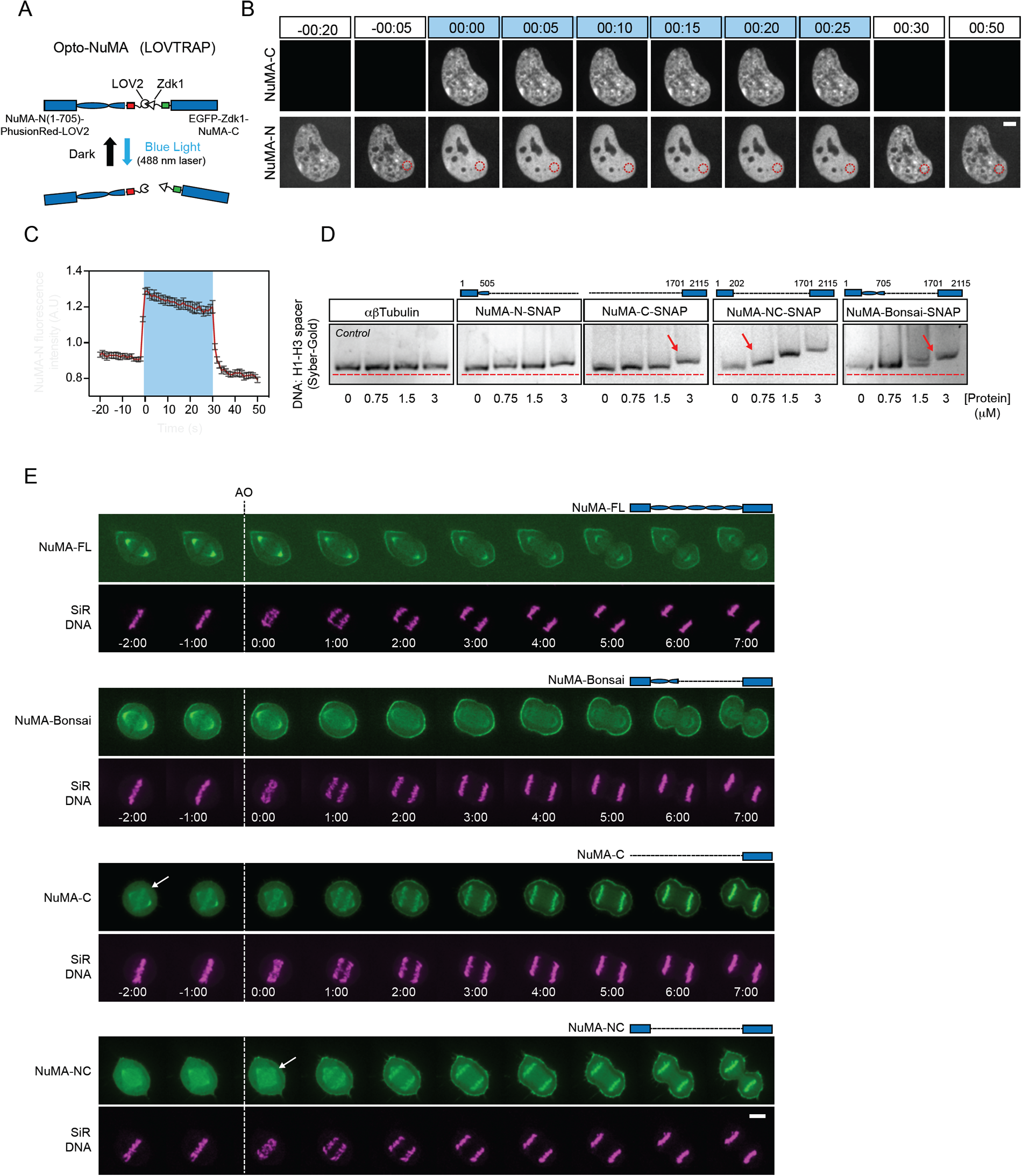
NuMA’s coiled-coil regulates its C-terminus’ chromatin binding over the cell cycle. **(A)** LOVTRAP light-induced dissociation system: in the dark, LOV2 (phototropin) binds to Zdk1, while blue light induces a conformational change in LOV2 that prevents binding to Zdk1. In the Opto-NuMA engineered protein NuMA-N(1-705)-PhusionRed-LOV2 + EGFP-Zdk1-NuMA-C, both NuMA ends are linked through LOVTRAP, and blue light dissociates them from each other. **(B)** Representative timelapse of an uninduced NuMA-KO RPE1 cell stably expressing Opto-NuMA in the nucleus before, during (0:00-0:25) and after illumination with blue (488 nm) light, showing NuMA-N but not NuMA-C becoming diffuse upon dissociation. EGFP-Zdk1-NuMA-C cannot be imagined without dissociating Opto-NuMA. Red circle corresponds to the area chosen for intensity analysis in (C). Time in min:s. Scale bar: 5 μm. **(C)** NuMA-N(1-705)-PhusionRed-LOV2 intensity in the nucleus before, during (blue box) and after illumination with blue light. Red trace = mean, black = SD. n=10 cells. **(D)** (Top) The schematics represent the NuMA constructs used in each gel, with amino acid numbers indicated. (Bottom) EMSA using 60 ng H1-H3 spacer DNA fragment and increasing concentrations of αβTubulin, NuMA-N-SNAP, NuMA-C-SNAP, NuMA-Bonsai-SNAP and NuMA-NC-SNAP. DNA was labeled with Sybr-Gold. Arrows indicate the minimum protein concentration for each protein with a noticeable DNA band shift. **(E)** Representative timelapse images of RPE1 cells transiently expressing NuMA-FL-EGFP and stably expressing NuMA-Bonsai-EGFP, EGFP-Zdk1-NuMA-C or NuMA-NC-EGFP and labelled with SiR-Hoechst (DNA), followed from metaphase through cytokinesis. NuMA-C and NuMA-NC localize to chromosomes at mitosis, while NuMA-Bonsai does not. Time in min:s, 0:00 corresponds to anaphase onset (AO, vertical dashed line). Arrows indicate when we detect different NuMA truncations on chromosomes. Scale bar: 5 μm.

Previous work suggested that full-length NuMA can directly bind DNA (Ludérus et al., 1994), but whether binding occurs through the C- or N-terminus or both, and the relative affinity of each for DNA, are not known. To address these questions and understand how NuMA binds DNA, we purified different SNAP-tagged NuMA truncation constructs (Fig. S4) and tested their ability to bind a DNA fragment *in vitro* using an electrophoretic mobility shift assay (EMSA) under stringent binding conditions (Fig. 4D). We observed that the construct that lacks the coiled-coil (NuMA-NC) bound DNA with the highest affinity, followed by NuMA-Bonsai (NuMA-NC with an additional 493 amino acid-long coiled-coil) and finally NuMA-C with the lowest detectable affinity. NuMA-N did not have detectable DNA affinity in our conditions, and negative control tubulin had no detectable affinity. Thus, NuMA directly binds DNA primarily through its C-terminus, and this binding could in principle underlie NuMA-C’s chromatin binding *in vivo* (Fig. 4A-C). Further, these *in vitro* observations suggest that NuMA’s DNA binding may be intra-molecularly regulated, as NuMA’s N-terminus increases its DNA binding and the coiled-coil appears to decrease it.

Given that NuMA-C binds chromatin in interphase (Fig. 4A-C) and DNA *in vitro* (Fig. 4D), we asked what prevents NuMA-C from binding chromosomes in mitosis. To answer this question, we mapped the localization of NuMA-FL, NuMA-Bonsai, NuMA-NC and NuMA-C through mitosis (Fig. 4E; Video 4), testing whether other parts of NuMA regulate NuMA-C’s chromatin binding, as suggested *in vitro* (Fig. 4D). Similar to endogenous NuMA (Fig. S3) and NuMA-FL, NuMA-Bonsai did not localize to chromosomes in mitosis. In contrast, NuMA-NC and NuMA-C were recruited to mitotic chromosomes as early as anaphase onset. Therefore, NuMA’s coiled-coil actively regulates NuMA’s localization during mitosis by preventing the premature binding of NuMA-C to chromosomes. NuMA is heavily phosphorylated by the mitotic kinase CDK1 at mitotic entry (Compton and Luo, 1995; Kotak et al., 2013; Yang et al., 1992). CDK1 activity negatively controls NuMA levels at the cell cortex during metaphase, and NuMA’s dephosphorylation at anaphase promotes the protein’s translocation to the cortex (Kiyomitsu and Cheeseman, 2013; Kotak et al., 2013). Premature RO3366-mediated CDK1 inhibition at metaphase did not change NuMA-Bonsai’s localization, while NuMA-C and NuMA-NC still bound to chromosomes (Fig. S5). This indicates that the coiled-coil regulates NuMA localization during mitosis, apparently doing so independent of CDK1 activity and may depend on other regulatory processes. Together, these findings indicate that the coiled-coil of NuMA acts as a central hub that not only drives the protein’s nuclear specific function and the stability of its assemblies (Fig. 3), but also when NuMA is allowed to bind chromosomes and which structures it acts on over the cell cycle.

### NuMA provides mechanical robustness to the nucleus

So far, we have shown that NuMA is essential for forming a single, round nucleus (Fig. 1-2), keeps the chromosome mass compact at mitotic exit (Fig. 2F), can form higher order nuclear assemblies (Fig. 3G; (Zeng et al., 1994)) and binds chromatin in a cell cycle-regulated manner (Fig. 4). Together, these findings indicate that NuMA plays a long-range structural role in forming the nucleus, and suggest how it may do so. This led us to hypothesize that NuMA also plays a structural role in maintaining the nucleus. To test for such a role, we compared nuclear morphologies and then mechanically perturbed the nuclei of cells stably expressing EGFP-Lap2β and mCherry-H2B with and without NuMA. If nuclei were more deformable without NuMA, this would indicate that NuMA plays a structural role in nuclear mechanics. Imaging in three dimensions, and at higher resolution than above experiments, revealed invaginations in the nuclear surface of NuMA KO cells that were undetectable in control cells (Fig. 5A). This suggested that nuclei are more deformable in NuMA KO cells. To more directly compare nuclear mechanics with and without NuMA, we confined and thereby compressed cells using a PDMS device (Guild et al., 2017; Le Berre et al., 2012) with 3 μm high pillars (Fig. 5B), and measured resulting changes in nuclear morphology. Under confinement, the nuclei of NuMA KO cells lost their nuclear invaginations as these ‘unfolded’ into a smooth nuclear surface similar to control cells (Fig. 5C). Further, nuclei without NuMA were more deformable under external mechanical force than control nuclei: while nuclei were thicker (i.e. higher in the confinement axis) prior to confinement in NuMA KO cells, they became thinner than control cells under confinement (Fig. 5D-F). Thus, NuMA provides mechanical stability to the nucleus, which could help maintain nuclear structure and function. Altogether, our work indicates that NuMA plays a spindle-independent, long-range structural role in both nuclear formation and mechanics.

**Figure 5.**
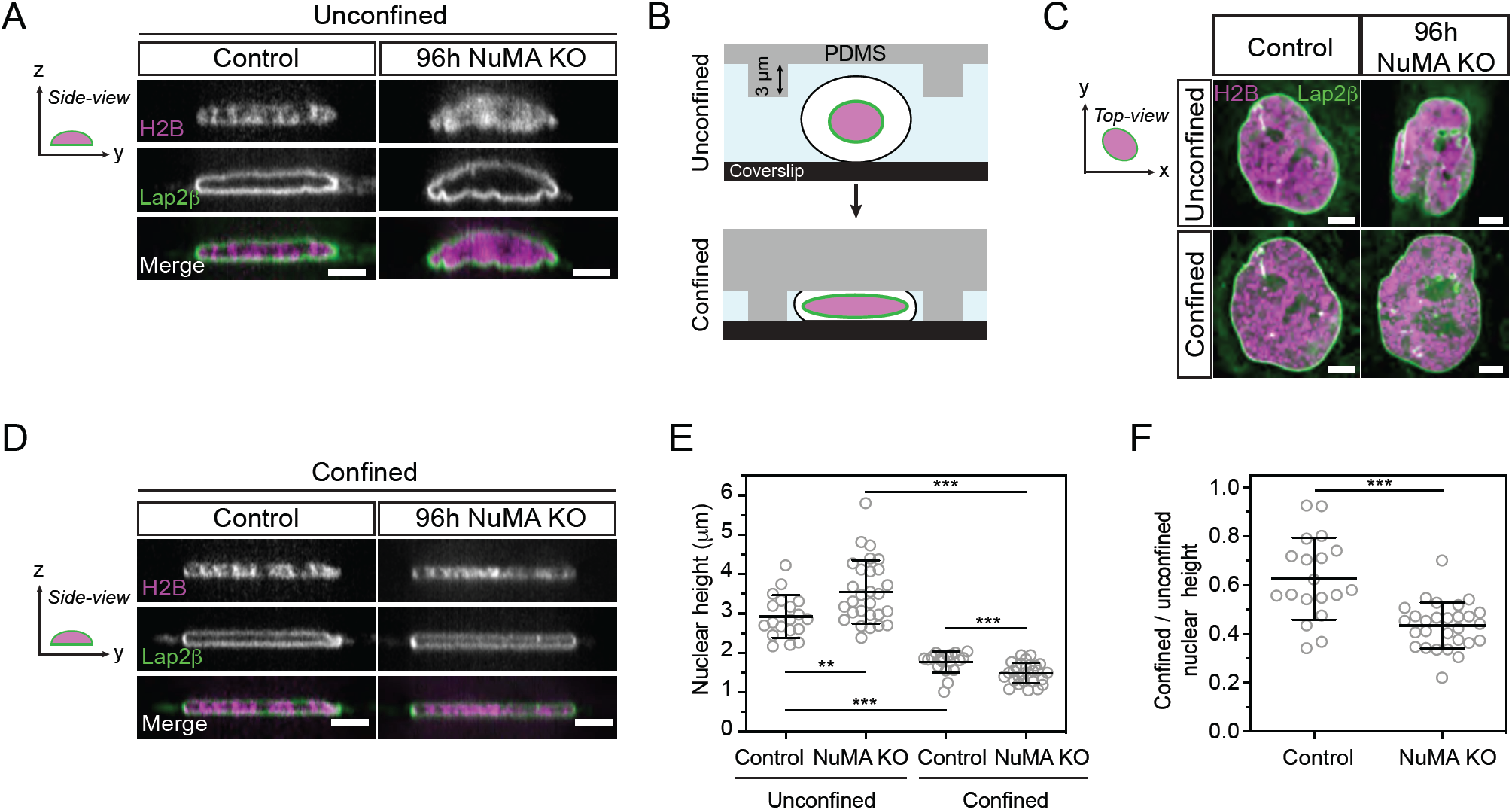
NuMA provides mechanical robustness to the nucleus. **(A)** Representative side (y-z) view images of a live control and 96h NuMA KO RPE1 cell expressing mCherry-H2B (magenta) and EGFP-Lap2β (green), with images acquired every 0.25 μm over 8 μm. Scale bar: 5 μm. **(B)** Schematic representation of cell confinement using a PDMS device (grey) with 3 μm high pillars. Nuclear (pink) compression occurs when pillars are brought down to contact the coverslip (black). **(C)** Top (x-y) view images of a live control and 96h NuMA KO RPE1 cells from (A), shown before (unconfined) and during confinement by the PDMS device depicted in (B). Scale bar: 5 μm. **(D)** Representative side (y-z) view images of a confined live control and NuMA KO RPE1 cell from (A) and (C). Scale bar: 5 μm. **(E-F)** Nuclear height of unconfined and confined control and NuMA KO cells (E), calculated based on EGFP-Lap2β localization, and unconfined nuclear height ratio in control and NuMA KO cells (F). Mean ± SD. n=28 (control) and 17 (NuMA KO) cells. Mann-Whitney test: **p < 0.01; ***p < 0.001.

## Discussion

Although many of the molecules required for nuclear formation are known, we still do not understand how they together give rise to a single, round and robust mammalian nucleus. Because mitotic errors can lead to abnormal nuclei, one challenge is to define when – at mitosis or nuclear formation – a given molecule acts, and what roles it plays at different times. A long standing example has been NuMA, a ∼ 200 nm-long coiled-coil protein (Harborth et al., 1995, 1999; Yang et al., 1992) essential for both mitosis (Compton and Luo, 1995; Kallajoki et al., 1991; Yang and Snyder, 1992) and nuclear integrity (Compton and Cleveland, 1993; Kallajoki et al., 1991, 1993). Here, we uncouple NuMA’s mitotic and nuclear functions. We show that NuMA has a spindle-independent role in building a single, round nucleus (Fig. 1), keeping the decondensing chromosome mass compact at nuclear formation (Fig. 2) and promoting the nucleus’ mechanical robustness (Fig. 5). Thus, NuMA plays a long-range structural role in both building and maintaining the nucleus, as it does in the spindle (Compton and Luo, 1995; Hueschen et al., 2019; Silk et al., 2009; Yang and Snyder, 1992). We find that NuMA’s coiled-coil acts as a hub: it is required for nuclear function, regulates NuMA’s mobility in the nucleus (Fig. 3) and its binding to chromosomes throughout the cell cycle (Fig. 4). As such, the coiled-coil compartmentalizes NuMA’s function to where and when it is needed: on chromosomes at interphase (when these are kept together) and off chromosomes at mitosis (when these are pulled apart). We propose a model whereby NuMA drives μm-scale cellular organization throughout the cell cycle, shaping and protecting the nucleus at interphase and the spindle at mitosis (Fig. 6).

**Figure 6.**
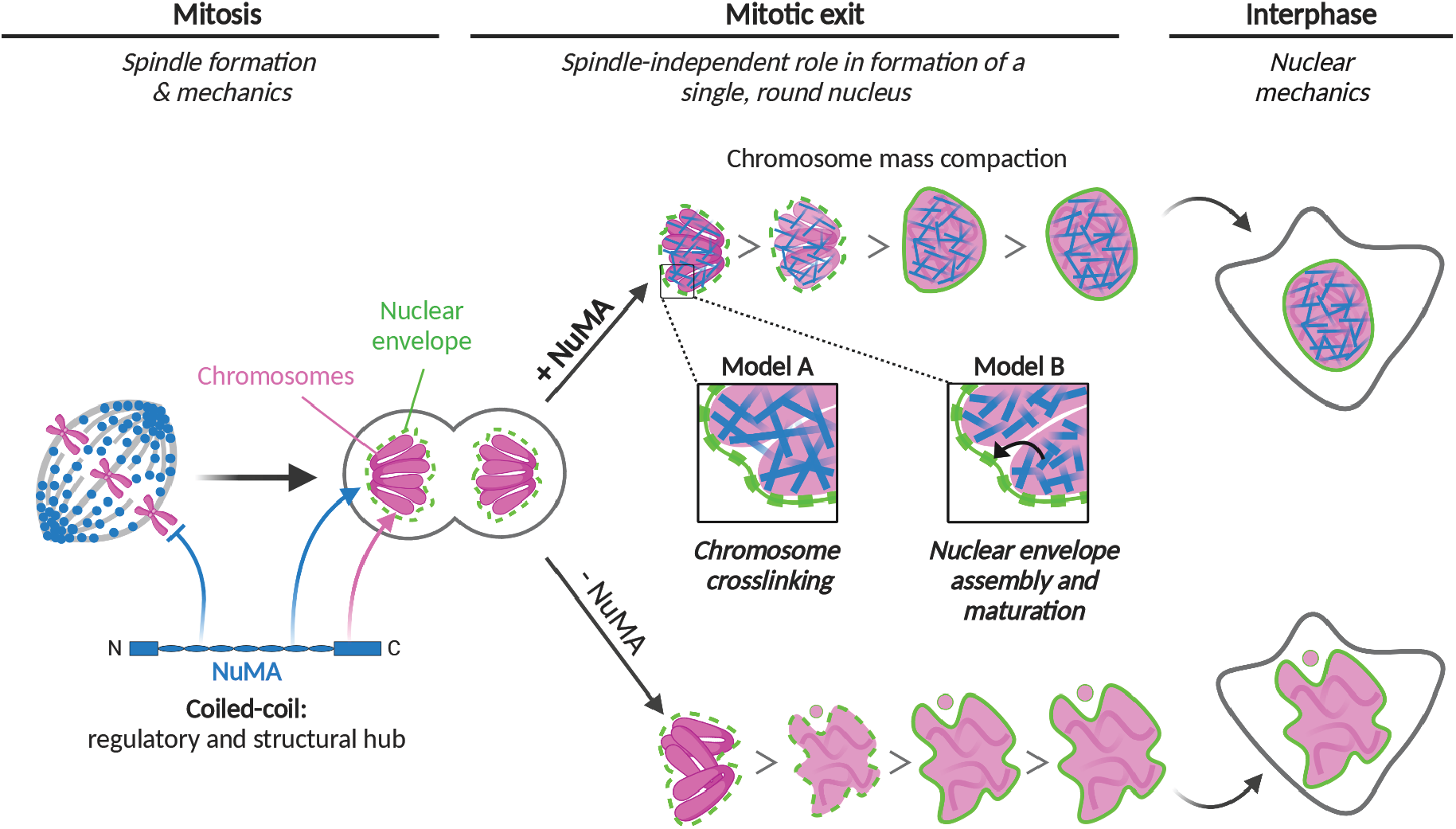
Model for NuMA’s role in nuclear formation and mechanics, and its long-range structural role over the cell cycle. NuMA (blue) plays a spindle-independent role in nuclear formation (“Mitotic exit”, center) and mechanics (“Interphase”, right). It keeps the chromosome (pink) mass compact at nuclear formation, and is essential to building a single, round and mechanically robust nucleus (“+NuMA”, top). Without NuMA (“-NuMA”, bottom), micronucleation and nuclear shape defects occur. We propose two models for how NuMA, whose C-terminus binds interphase chromosomes (pink arrow), performs its nuclear function. To promote nuclear formation and mechanics, NuMA could crosslink chromosomes (Model A, blue filaments) or regulate nuclear envelope (green) assembly and maturation (Model B, black arrow), either directly or indirectly. At “Mitosis” (left), NuMA plays a critical role in spindle formation and mechanics and its coiled-coil prevents it from binding chromosomes, when these must but segregated instead of kept together. At mitotic exit and interphase, the coiled-coil drives NuMA’s nuclear dynamics and function (blue arrows). As such, we propose that NuMA’s coiled-coil acts as a regulatory and structural hub to control its function in space and time. Altogether, NuMA is essential to the formation and mechanics of two of the cell’s largest structures, the spindle and the nucleus, protecting genome integrity across the cell cycle.

### NuMA’s structure, regulation and function in the nucleus

Our current understanding of NuMA’s spindle function and its underlying basis much surpasses that of its nuclear function. Here, our work raises the question of how NuMA’s structure can give rise to its function in nuclear formation (Fig. 1-2) and mechanics (Fig. 5). While most of NuMA’s coiled-coil is dispensable for building the spindle (Forth et al., 2014; Hueschen et al., 2017), it is essential for building a single, round nucleus (Fig. 3B-C; S2C). Additionally, the coiled-coil can stabilize nuclear NuMA (Fig. 3D-F), and overexpressing full-length NuMA, but not coiled-coil truncations, induces the formation of stable filament-like nuclear structures (Fig. 3G). Further, we know that NuMA can, when overexpressed, assemble into a three-dimensional network whose lattice size is determined by its coiled-coil’s length (Gueth-Hallonet et al., 1998; Harborth et al., 1999). We also know that NuMA self-assembles at the cell cortex to position the spindle using its coiled-coil and C-terminus (Okumura et al., 2018). Together, these observations are consistent with a model where NuMA assembles into a network that shapes nuclear morphology and mechanics (Fig. 1-2, 5), with its coiled-coiled key to both assembly and function. Since NuMA-C binds DNA and chromosomes (Fig. 4), such a network could in principle help crosslink DNA within and between chromosomes. NuMA’s long structure and ability to self-assemble in principle make it well-suited to bridge large distances. Determining to what extent nuclear NuMA self-assembles, and the structure of this assembly, will be essential to understanding how NuMA’s structure gives rise to its dynamics (Fig. 3D-F) and architectural and mechanical functions (Fig. 1-2, 5) in the nucleus. Uncovering NuMA’s nuclear functions may in turn help understand why its expression levels vary dramatically across cell types and states (Merdes and Cleveland, 1998).

While NuMA-C binds DNA *in vitro* (Fig. 4D) and appears to bind chromosomes in the nucleus (Fig. 4A-B), full-length NuMA does not bind chromosomes at mitosis. As such, its binding must be regulated. Here, we show that a small portion of NuMA’s coiled-coil is sufficient to prevent NuMA-C from binding chromosomes at mitosis (Fig. 4E), and sufficient to decrease NuMA’s DNA binding affinity *in vitro* (Fig. 4D). Preventing NuMA from binding chromosomes at mitosis could prevent it from keeping chromosomes together when they must segregate or, since NuMA recruits dynein (Hueschen et al., 2017), from recruiting dynein-based forces there. However, we find that prematurely localizing NuMA-NC on chromosomes at mitosis does not lead to detectable mitotic defects (Fig. 4E). This could be because NuMA-NC lacks the coiled-coil needed for NuMA to keep chromosomes together (Fig. 3B-C; S2C) and likely necessary to activate dynein motility (Hueschen et al., 2017; Reck-Peterson et al., 2018), or because its function on chromosomes is cell-cycle regulated. We propose NuMA’s coiled-coil plays a central role throughout the cell cycle, acting as a structural (Fig. 3B-C; S2C) and regulatory (Fig. 3D-F, 4E) hub for NuMA’s function on chromosomes. How the cell regulates this hub, such that NuMA binds chromosomes at mitotic exit and dissociates at mitotic entry, remains an open question. Answering it will require defining how NuMA’s C-terminus binds DNA and chromosomes, and will be key to understanding how the same protein performs such different physical roles in interphase and mitosis. Notably, NuMA’s C-terminus also targets NuMA in the spindle, specifically to microtubule minus-ends (Hueschen et al., 2017).

### Models for NuMA’s role in nuclear formation and mechanics

What role NuMA plays in nuclear formation (Fig. 1-2) and mechanics (Fig. 5) is unclear. One model is that NuMA directly contributes to the organization and mechanics of the chromosome mass at mitotic exit (Fig. 6, model A). Indeed, NuMA-C binds DNA *in vitro* (Fig. 4D) and chromosomes *in vivo* (Fig. 4A-C), NuMA keeps the mass of decondensing chromosomes compact at mitotic exit (Fig. 2F), and it localizes throughout the entire nucleus ((Compton et al., 1992; Yang et al., 1992); Fig. S3), where it can interact with chromosomes. Further, NuMA influences higher order chromatin organization (Abad et al., 2007; Kivinen et al., 2010) and binds chromatin remodelers (Vidi et al., 2014), and we know that chromatin compaction impacts the shape and robustness of the nucleus (Stephens et al., 2019). If NuMA crosslinked DNA and chromosomes, for example, its import into the nucleus as it is forming and chromatin starts decondensing (Schooley et al., 2012) could keep the chromosome mass more compact and rigid. This could make it harder for the assembling nuclear envelope to penetrate into the chromosome mass, thus decreasing multinucleation and abnormally shaped nuclei (Fig. 1-2), and provide mechanical robustness to the nucleus (Fig. 5). In this model, NuMA plays a nuclear formation role with similarities to BAF’s, with BAF localizing at the chromosome mass’ rim (Samwer et al., 2017) and NuMA through the entire volume of the nucleus. Finally, we note that NuMA could also organize chromosomes at mitotic exit by keeping them in a tight focus around spindle poles (Cleveland, 1995; Compton and Cleveland, 1993), given NuMA’s pole focusing role. However, such a role cannot explain our observations that NuMA plays a role in nuclear formation after a spindle-less mitosis (Fig. 1-2).

Alternatively, and not mutually exclusive, NuMA’s binding to chromosomes during nuclear formation could promote assembly and maturation of the nuclear envelope and its components (Fig. 6, model B; (LaJoie and Ullman, 2017; Ungricht and Kutay, 2017)) or mediate the tethering of chromatin to the nuclear envelope (Schreiner et al., 2015). These, when compromised, could lead to defects in nuclear morphology (Fig. 1-2) and mechanics (Fig. 5) (Davidson and Lammerding, 2014; Webster et al., 2009). Incidentally, NuMA-C-coated beads mixed with *Xenopus* extracts recruit membranes, suggesting that NuMA could contribute to nuclear envelope assembly (Lu et al., 2012). Though NuMA is not enriched at the nuclear rim (Fig. 1A, S3), we cannot exclude that it regulates the nuclear envelope and as such indirectly impacts chromosome organization.

Looking forward, testing these and other models will be critical to understanding NuMA’s physical role in building and maintaining the nucleus. Towards this goal, it will be important to determine whether NuMA can crosslink chromatin *in vitro* and *in vivo*, and whether and how it influences the nuclear envelope. Tools to acutely inhibit NuMA will be essential to separating its roles in nuclear formation versus maintenance. Defining NuMA’s physical role in the nucleus, and with which molecular players it acts, will be key to understanding its relationship to other mechanisms regulating nuclear formation and maintenance. More broadly, doing so will provide insight into how NuMA is repurposed and regulated to protect both the nucleus and spindle, preserving genome integrity across the cell cycle.

## Acknowledgements

We thank Christina Hueschen for inducible RPE1 NuMA-KO cells; Daniel Gerlich for reagents; Jeffrey van Haren and Torsten Wittmann for LOVTRAP plasmids and discussions; Abby Buchwalter, Daniel Gerlich and Megan King for discussions; Joël Lemière and Arthur Molines for help with FRAP analysis and discussions; Amanda Jack for advice on DNA binding experiments; Delaine Larson and Kari Herrington at the UCSF Nikon Imaging Center for assistance with FRAP experiments; Abby Buchwalter, Christina Hueschen and Arthur Molines for critical reading of the manuscript; the Fred Chang laboratory for discussions and the Dumont laboratory for discussions and critical reading of the manuscript. We thank Sachin Kotak for generously sharing findings prior to publication. The FRAP data for this study were acquired at the UCSF Nikon Imaging Center with instruments obtained using funding from the NIH (5R35GM118119), the UCSF Program for Breakthrough Biomedical Research funded in part by the Sandler Foundation, the UCSF Research Resource Fund Award, and HHMI. We acknowledge the UCSF Parnassus Flow Cytometry Core supported in part by the NIH P30 DK063720 grant and the NIH S10 Instrumentation Grant S10 1S10OD021822-01 for assistance. This work was funded by NIH DP2GM119177 (S.D.), NIH R01GM134132 (S.D.), the Rita Allen Foundation (S.D.), the NSF Center for Cellular Construction 1548297 (S.D.), NSF CAREER 1554139 (S.D.), NSF MCB-1617028 (A.Y.) and a HFSP Long Term Postdoctoral Fellowship (A.S.-M.).

The authors declare no competing financial interests.

## Author contributions

Andrea Serra-Marques, Conceptualization, Methodology, Software, Validation, Formal Analysis, Investigation, Resources, Data Curation, Writing – Original Draft Preparation, Writing – Review & Editing, Visualization, Supervision, Funding Acquisition; Ronja Houtekamer, Conceptualization, Methodology, Validation, Formal Analysis, Investigation, Data Curation, Writing – Review & Editing, Visualization; Dorine Hintzen, Conceptualization, Methodology, Validation, Investigation, Writing – Review & Editing; John T. Canty, Methodology, Validation, Investigation, Data Curation, Writing – Review & Editing, Visualization; Ahmet Yildiz, Methodology, Writing – Review & Editing, Supervision, Funding Acquisition; Sophie Dumont, Conceptualization, Methodology, Resources, Writing – Review & Editing, Supervision, Funding Acquisition.

## Methods

### Cell culture and transfection

Tet-on inducible CRISPR/Cas9 NuMA KO RPE1 cells (hereafter “NuMA KO RPE1 cells”) (Hueschen et al., 2017) were cultured in DMEM/F12 with GlutaMAX (10565018; Thermo Fisher) supplemented with 10% tetracycline-screened fetal bovine serum (SH30070.03T; Hyclone Labs). Cells were maintained at 37 °C and 5 % CO_2_. NuMA was depleted by inducing spCas9 expression with 1 μg/ml Doxycycline hyclate in Tet-on inducible CRISPR/Cas9 RPE1 sgNuMA cells for 96 h before each experiment, unless other induction times are indicated. For transient expression experiments (Fig. 3B, 3D-G (NuMA-FL-EGFP only), 4E (NuMA-FL-EGFP only), S3A-B, S3E-F (NuMA-FL-EGFP only), inducible NuMA KO RPE1 cells were transfected with DNA using ViaFect (E4981; Promega) according to manufacturer’s instructions 48 h before imaging. For rescue experiments (Fig. 3B, S3A-B), cells were transfected 72 h before imaging. Cell lines were not STR-profiled for authentication. All cell lines tested negative for mycoplasma.

### Cell line generation and lentiviral production

To generate a cell line stably expressing mCherry-H2B and EGFP-Lap2β, NuMA KO RPE1 cells were infected with mCherry-H2B and EGFP-Lap2β lentiviruses (Fig. 2, 5). Viruses were made in HEK293T cells from a pLenti6/V5-DEST plasmid (Invitrogen) containing mCherry-H2B (gift from Torsten Wittmann) or EGFP-Lap2β and lentiviral packaging plasmids using calcium phosphate transfection. The media of transfected HEK293T cells was replaced after 24 h and virus-containing media was collected 48 h after transfection. NuMA KO RPE1 cells were infected with virus-containing media supplemented with 10 μg/ml polybrene and selected with 2.5 μg/ml blasticidin. A polyclonal stable line with low expression of mCherry-H2B and EGFP-Lap2β was further selected by FACS sorting and used herein.

To generate the Opto-NuMA cell line (Fig. 4B-C) NuMA KO RPE1 cells were infected with NuMA-N-PhusionRed-LOV2 and EGFP-Zdk1-NuMA-C using retroviruses made in HEK293T-GP cells transfected with calcium phosphate, as described above. A polyclonal stable line was established after FACS sorting.

To generate cell lines stably expressing NuMA truncations, the above protocol was used to infect NuMA KO RPE1 cells. We made the following cell lines: monoclonal NuMA-Bonsai-EGFP (Fig. 3D-F, 4E, S2B-D, S5), EGFP-Zdk1-NuMA-C (Fig. 4E, S5) and NuMA-NC-EGFP (Fig. 3D-F, 4E, S5). In the NuMA-Bonsai-EGFP line, exogenous NuMA is expressed at higher levels than endogenous NuMA (Fig. S2B).

### Drug and dye treatments

Cells were treated with 664 nM nocodazole (M1404; Sigma-Aldrich) to depolymerize spindle microtubules and with 320 nM reversine (R3904; Sigma-Aldrich) to inhibit MPS1 and silence the spindle assembly checkpoint (Fig. 1, S1A-C). To avoid the confounding negative effect of a too-short mitosis on successful nuclear formation, we adjusted reversine concentrations in control cells (1 μM) so that NuMA KO cells spent at least as much time in spindle-less mitosis as control cells (Fig. 2, S1D-E). Nocodazole and reversine were added 10-30 min before live imaging. Cells were treated with 9 μM RO-3306 to acutely inhibit CDK1. To visualize DNA by live imaging in cells not expressing mCherry-H2B (Fig. 4E; S5), 500 nM SiR-DNA dye (SC007; Cytoskeleton) was added 30 min before imaging.

### Plasmids

To make the mCherry-H2B and EGFP-Lap2β cell line, the EGFP-Lap2β coding sequence was amplified from a EGFP-Lap2β plasmid (gift from Daniel Gerlich) (Samwer et al., 2017), and inserted into the lentiviral vector pLenti6/V5-DEST plasmid (Invitrogen, gift from Torsten Wittman). The lentiviral mCherry-H2B plasmid was a gift from Torsten Wittman (Pemble et al., 2017).

To generate the Opto-NuMA cell line, NuMA-N-PhusionRed-LOV2 was cloned into a CMV promoter plasmid as follows: NuMA-N^1-705^ was amplified from NuMA-Bonsai^1-705+1701-2115^-EGFP (Hueschen et al., 2017) and PhusionRed was amplified from pmKate2.7C-N1 (the Michael Davidson Fluorescent Protein Collection, UCSF Nikon Imaging Center). These were subsequently cloned into a CMV promoter plasmid containing the fast dissociation LOV2 domain (EB1-N-LZ-LOV2 described in van Haren et al., 2018), by replacing the EB1-N-LZ sequence with the NuMA-N-PhusionRed sequence. EGFP-Zdk1-NuMA-C was cloned into the CMV promoter plasmid mCherry-Zdk1-EB1-C (van Haren et al., 2018) as follows: mCherry was replaced by EGFP amplified from pEGFP-C1 (Clonetech, Takara Bio) and EB1-C was replaced by NuMA-C^1701-2115^, which was amplified from NuMA-Bonsai-EGFP (Hueschen et al., 2017). C) As a final step, NuMA-N-PhusionRed-LOV2 and EGFP-Zdk1-NuMA-C were subcloned into a Moloney murine leukemia retroviral plasmid (gift from Iain Cheeseman; pIC291, Addgene #51403 (Welburn et al., 2009)). For this, pIC291 was digested with BamHI to remove EGFP-TEV-S-Ska1, and NuMA-N-PhusionRed-LOV2 and EGFP-Zdk1-NuMA-C were amplified and inserted.

To generate cell lines stably expressing NuMA truncations, NuMA-Bonsai^1-705+1701-2115^-EGFP was amplified from NuMA-Bonsai^1-705+1701-2115^-EGFP (Hueschen et al., 2017), EGFP-Zdk1-NuMA-C was made as described above, and NuMA-NC^1-212+1701-2115^-EGFP was made from assembling both NuMA-N^1-212^ and NuMA-C^1701-2115^ from NuMA-Bonsai^1-705+1701-2115^-EGFP (Hueschen et al., 2017), then these were each inserted into the lentiviral vector pLenti6/V5-DEST plasmid (Invitrogen, gift from Torsten Wittmann). These three truncation constructs are resistant to the NuMA sgRNA, as described in (Hueschen et al., 2017).

NuMA-FL-EGFP and NuMA-Bonsai-EGFP used for transient transfection are from NuMA-Bonsai^1-705+1701-2115^-EGFP (Hueschen et al., 2017); for reference, the NuMA-Bonsai-EGFP herein is called NuMA-N-C-EGFP in (Hueschen et al., 2017). The NuMA-NC-EGFP construct used for transient transfection is the same used to generate the stable cell line. Notably, NuMA-FL-EGFP, NuMA-Bonsai-EGFP and NuMA-NC-EGFP plasmids are resistant to the NuMA sgRNA, as described in (Hueschen et al., 2017).

To express and purify proteins for EMSA, the SNAP-tagged NuMA proteins were cloned as follows: NuMA-N^1-505^, NuMA-C^1701-2115^, NuMA-Bonsai^1-705+1701-2115^ and NuMA-NC^1-212+1701-2115^ were amplified from NuMA-FL-EGFP (Hueschen et al., 2017) and inserted into an Sf9 expression vector (pACEBac1). αβTubulin was purified from porcine brains (Hyman et al., 1991).

### Western blotting

Cells were plated in 6-well plates (08-772-1B; Fisher Scientific) in parallel with setting up an imaging experiment. Media was removed and cells were collected at the indicated time after induction of NuMA KO. Cells were subsequently lysed in cold RIPA buffer, supplemented with protease inhibitor cocktail (11836153001; Roche), incubated on ice for 30 min and protein extracts collected after 30 min of centrifugation at 4 °C. Protein concentrations were measured using a Bradford assay kit (Bio-Rad) based on absorbance (A595 nm), and samples were prepared at equal concentrations in NuPage Sample buffer and NuPage Sample reduction agent (Invitrogen). Proteins were denatured for 10 min at 95 °C, separated on a 3-8 % Tris-Acetate gel (Invitrogen) by SDS-PAGE, and transferred to a nitrocellulose membrane by Western blotting using a Bio-Rad electrophoresis system. Membranes were blocked with 4 % milk in TBS-T for 1 h at room temperature, probed with rabbit anti-NuMA (1:1000; ab97585; Abcam; Fig. S2A-C), rabbit anti-NuMA (1:300; NB500-174; Novus Biologicals; Fig. S1A-D) and mouse anti-α-Tubulin (1:5000; T6199; Sigma-Aldrich) primary antibodies in 1 % milk in TBS-T at 4 °C overnight, washed with TBS-T, and incubated with anti-rabbit and anti-mouse HRP-conjugated secondary antibodies (1:10000; sc-2357 and sc-2005; Santa Cruz Biotechnologies) for 1 h at room temperature. Proteins were detected with the ImageQuant LAS4000 using ECL (Thermo Fisher) based chemiluminescence.

### Immunofluorescence

Cells were plated on 25 mm glass coverslips coated with 1mg/ml poly-L-lysine. Cells were fixed with 95 % (v/v) methanol containing 5 mM EGTA for 2 min at −20 °C, washed with 0.05 % Triton-X-100 in Tris-Buffered Saline (TBS-T), and blocked with 2 % Bovine Serum Albumin (BSA) in TBS-T for 1 h at room temperature. Primary and secondary antibodies were diluted in 2 % BSA in TBS-T and incubated with cells for 1 h at room temperature. DNA was labeled with Hoechst 33342 (B2261; Sigma). Cells were mounted in ProLongGold Antifade (P36934; Thermo Fisher). Cells were imaged using a spinning disk confocal inverted or an epifluorescence microscope as described in the following section. The following antibodies were used: rabbit anti-NuMA (1:300; NB500-174; Novus Biologicals; Fig. 1, S1A-E, S3), rabbit anti-NuMA (1:130; ab97585; Abcam; Fig. 3B-C, S2A,C), anti-GFP conjugated to Atto488 (1:100; ChromoTek gba-488; Fig. 3B-C, S2A,C), anti-mouse secondary antibodies (1:400) conjugated to Alexa Fluor 488 (A11001; Invitrogen), anti-rabbit secondary antibodies (1:400) conjugated to Alexa Fluor 647 (A21244; Invitrogen).

### Imaging

NuMA KO RPE1 cells were plated on #1.5 glass-bottom 35 mm MatTek dishes coated with poly-D-lysine (MatTek Corporation). Imaging was performed on a spinning-disk confocal (CSU-X1; Yokogawa) inverted microscope (Eclipse Ti-E, Nikon instruments) with a perfect focus system (Nikon). Experiments were performed with a Di01-T405/488/568/647 head dichroic (Semrock), along with 405 nm (100-mW), 488 nm (150 mW), 561 nm (150-mW) and 642 nm (100 mW) diode lasers, and emission filters ET455/50M, ET525/50M, ET630/75M and ET705/72 (Chroma). Cells were imaged with a Zyla 4.2 sCMOS camera (Andor Technology) via Metamorph (7.10.3, MDS Analytical Technologies) by fluorescence (50-100 ms exposures) with a 60 × 1.4 Ph3 oil objective yielding 109 nm/pixel at bin = 1. For long-term live imaging quantifications (Fig. 2), a 20X 0.5 Ph1 air objective yielding 329 nm/pixel at bin = 1 was used. For long-term live imaging (Fig. 2), images were acquired every 4 min over 11 z-planes spaced 2 μm apart to account for focal plane changes over long periods. All other live images were acquired in a single plane, every 1-60 s. For fixed cell imaging, images were acquired over 20 z-planes spaced 0.3 μm apart. For the confinement experiments (Fig. 5), images were acquired over 32 z-planes spaced 0.25 μm apart. Fixed cell imaging in Figure S4 was performed on a Zeiss AxioPlan2 epifluorescence microscope. Imaging for the FRAP experiments is described in the FRAP section.

For long-term live imaging in Figure 2, cells were treated with nocodazole and reversine at 72 h post KO induction, and imaged for 16-18 h. NuMA depletion at the end of the live imaging has occurred for 88-90h, but for simplicity we refer to it as “96 h”. At 88-90 h of induction, NuMA is expected to be nearly fully depleted (Fig. S1A).

### SNAP-tagged protein expression and purification

NuMA-N^1-505^-SNAP, NuMA-Bonsai-SNAP, and NuMA-C^1701-2115^-SNAP were purified from baculovirus-infected Sf9 cells. Briefly, cell pellets were resuspended in lysis buffer (50 mM HEPES pH 7.4, 300 mM NaCl, 10% Glycerol, 1mM DTT, 2 mM PMSF) supplemented with 1x protease inhibitor cocktail (Roche cOmplete Protease Inhibitor Cocktail) and lysed using a Dounce homogenizer (20 strokes with loose plunger followed by 20 strokes tight plunger). The lysate was clarified by centrifugation (65,000 rpm for 45 min) and incubated with IgG Sepharose (GE Healthcare) for 2 h at 12 °C, applied to a gravity flow column, and washed extensively with wash buffer (same as lysis buffer without PMSF or protease inhibitor cocktail). Protein labelling was performed by concentrating bead slurry to 5 ml, followed by the addition of 5 nmol of BG-LD555 dye (Lumidyne) then incubation (1 h at 12 °C). The slurry was re-added to a gravity flow column and washed extensively in wash buffer. The protein-bead complexes were then treated with TEV protease at 12 °C overnight. The mixture was then centrifuged (4000 rpm for 5 min) and the supernatant was retrieved. The mixture was then concentrated using an Amicon Ultra-0.5 ml 50k spin column (EMD Millipore). Protein concentration was determined using the Nanodrop 1000.

### Electrophoretic Mobility Shift Assay (EMSA)

Binding reactions were performed by mixing NuMA truncation proteins with 60 ng H1-H3 spacer DNA (Ludérus et al., 1994) followed by incubation on ice for 1 h. Binding buffer consisted of 20 mM HEPES pH 7.9, 100 mM NaCl, 50 nM KCl, 1 mM MgCl2, 1 mM DTT, 5 % glycerol, 0.1 mg/mL BSA, 0.1 % NP-40. After incubation, binding reactions were then loaded onto a 2 % Tris-Glycine agarose gel which was run for 90 min at 90 V. The gel was then post-stained with 1x Sybr-Gold (S11494; Thermo Fisher Scientific) for 20 min followed by UV imaging to visualize DNA.

### Fluorescence Recovery After Photobleaching (FRAP)

NuMA KO RPE1 cells transiently overexpressing NuMA-FL-EGFP, or stably expressing NuMA-Bonsai-EGFP, NuMA-NC-EGFP or EGFP-Zdk1-NuMA-C were plated on #1.5 glass bottom 35 mm MatTek dishes coated with Poly-L-lysine (MatTek Corporation) for 48 h. FRAP was performed on a OMX-SR microscope (GE Health Care), equipped with a PCO Edge 5.5 cMOS cameras (PCO), a four line laser launch 405/488/568/640 nm (Toptica), and live cell chamber (GE Health Care) at 37 °C with 5 % CO_2_. All images were acquired through a Plan ApoN 60x/1.42 NA (Olympus) oil immersion objective with Laser liquid 1.518 (Cargile). GFP was imaged with the 488 nm laser, with a 528/48M emission filter in place. Photobleaching was performed on a 2 µm diameter circular region in the nucleus. After defining the region of interest, a 405 laser was used at 40 % power for 150 ms to bleach the region. Images were acquired at 250 ms intervals for a total of 39 s; five baseline images were captured 1 s before bleaching and a timelapse of 39 s after the bleaching event.

### NuMA optogenetic control

NuMA KO RPE1 cells, non-induced and stably expressing NuMA-N-PhusionRed-LOV2 and EGFP-Zdk1-NuMA-C were imaged with a 561 nm laser (1 s interval) to visualize NuMA-N-PhusionRed-LOV2. The 488 nm laser was subsequently turned on (1s interval) to simultaneously activate the LOV2 domain and visualize EGFP-Zdk1-NuMA-C. The 488 nm laser was turned off after 30 s to end activation of the LOV2 domain while the 561 nm was used to continue imaging the behavior of NuMA-N-PhusionRed-LOV2.

### Cell confinement

Cells were plated on #1.5 glass-bottom 35 mm MatTek dishes coated with poly-D-lysine (MatTek Corporation). A polydimethylsiloxane (PDMS)-based suction cup device that contains a 10-mm-diameter glass coverslip with PDMS-based pillar structures (3 μm height; 200 μm diameter; spacing 700 μm to center) was generated (Guild et al., 2017; Le Berre et al., 2012). The device was attached to a 1 ml syringe, placed on top of a glass-bottom 35 mm dish, and sealed by pulling the syringe to apply negative pressure. After imaging 0.3 μm spaced z-planes of unconfined cells and initiating live acquisition, additional negative pressure was created to lower the pillared coverslips onto cells. At maximum confinement, the pillars prevent further cell compression. The negative pressure was maintained while z-stacks of compressed cells were acquired. Confinement was applied gradually over a period of about 1 min and sustained for 5 min.

### Data analysis

#### Nuclear phenotype classification (Fig. 1-3)

Observed nuclear phenotypes were qualitatively categorized into (1) single, round (absence of clear lobules or deep invaginations) nucleus, (2) nucleus with 1-3 micronuclei, (3) fragmented nucleus (> 3 micronuclei or multiple larger nuclei), and (4) atypical shape nucleus (cells with a single, abnormally shaped nucleus) based on DNA signal (Hoechst or mCherry-H2B). To determine percentages of these phenotypes before and after spindle-less mitosis (Fig. 2B), the phenotype was analyzed based on mCherry-H2B signal for all cells at t=0 (about 10 min after nocodazole and reversine addition) and at about t=16h.

#### Nuclear solidity and nuclear fluorescence intensity analysis (Fig. 1, S1-2)

A threshold of Hoechst or mCherry-H2B intensity was calculated and used to extract the outline of the nucleus using a particle analysis plugin from ImageJ. Based on this outline, the area and convex hull area were extracted.

Solidity was calculated as the ratio of the area of the nucleus to the area of a convex hull (Fig. 1D-E, S1D-E). NuMA and EGFP-tagged NuMA proteins mean intensities were measured in the nucleus based on a threshold of mCherry-H2B signal, as described above. The mean background intensity from three independent regions was subtracted. For each experiment, all values were normalized to the mean value of control (non-induced) cells (Fig. S1A, S1E).

#### Mother-daughter cell analysis (Fig. 2E)

Individual cells were followed and only cells that entered and exited spindle-less mitosis during image acquisition were included in the analysis. Phenotypes were determined before mitotic entry (mother cells) and after mitotic exit (daughter cells) for all cells. Only cells that had a single and round nucleus at mitotic entry were analyzed.

#### Nuclear expansion analysis (Fig. 2F)

Control and NuMA KO cells stably expressing mCherry-H2B and EGFP-Lap2β treated with nocodazole and reversine were randomly selected for analysis. NER (nuclear envelope reformation) (time=0) was determined by the first appearance of EGFP-Lap2β around condensed mitotic chromosomes and the area of the chromosome mass was calculated for the subsequent 132 min. Area was calculated based on a threshold of mCherry-H2B intensity using a particle analysis plugin from ImageJ. All thresholds were checked and manually corrected when automated threshold was not accurate before areas were calculated.

#### FRAP analysis (Fig. 3D-G, S2E-F)

The mean fluorescence intensity value in the bleached and non-bleached area, as well as background, were measured using a custom macro in ImageJ written by Arthur Molines. The signal was corrected for background and photobleaching and normalized 0 to 1, with 0 corresponding to t=0 and 1 to t=39s. Recovery measurements were quantified by fitting normalized fluorescence intensities of bleached areas to a double exponential for EGFP-tagged NuMA constructs using GraphPad Prism. Cells with an average intensity 3.5 standard deviations above the NuMA-FL-EGFP mean intensity were excluded. The recovery halftimes were calculated from double exponential fits (FL: R^2^= 0.87; Bonsai: R^2^=0.91; NC: R^2^=0.87).

#### NuMA-N photo-dissociation analysis (Fig. 4B-C)

NuMA-N intensity in the nucleus was measured on a 2.2 μm diameter circular region before, during and after illumination with a 488 nm laser. The signal was corrected for background and normalized to average intensity.

#### Nuclear confinement analysis (Fig. 5E-F)

To determine nuclear height before and after compression, orthogonal XZ and YZ views were generated from z-stacks of live cells (0.25 μm step size; 8 μm range), a line was drawn from top to bottom of the nucleus (based on EGFP-Lap2β signal) and its length was determined in ImageJ.

### Statistics

All data are expressed as average ± standard deviation (SD) or standard error of the mean (SEM), as indicated in the figures. All statistical analyses were carried out using Prism software (Graphpad). Data were tested for normality using the D’Agostino-Pearson normality test. P-values were calculated using unpaired t-tests or Mann-Whitney U tests as indicated in the figure legends. For the contingency tables of Fig. 1C, E, F, 2B, 2C and 3C, the Fisher’s exact test was applied. All p-values were two-tailed. Quoted n’s are described in more detail in figure legends, and in general refer to individual measurements (individual cells, nuclei).

### Image presentation

Timelapse images show single planes from spinning disk confocal imaging, except Figures 2A and 2D which show max intensity projections (20 μm total z distance) of spinning disk confocal z-stacks. Immunofluorescence images of nuclei and mitotic spindles (Fig. 1B, S1B, S1D) show maximum intensity projections (20 - 30 μm total z distance) of spinning disk confocal z-stacks. All Images and movies were formatted using ImageJ.

### Video preparation

Videos 1A and 1B show a maximum intensity projection from a spinning disk confocal z-stack. Video 2 shows a single z-slice from an OMX-SR microscope. Videos 3-4 show a single z-slice from a spinning disk confocal microscope. Videos were prepared for publication using ImageJ and set to play at 10 (Video 1A, 1B, 2 and 4) or 20 frames per second (Video 3).

## Supplemental Figure legends

**Figure S1.**
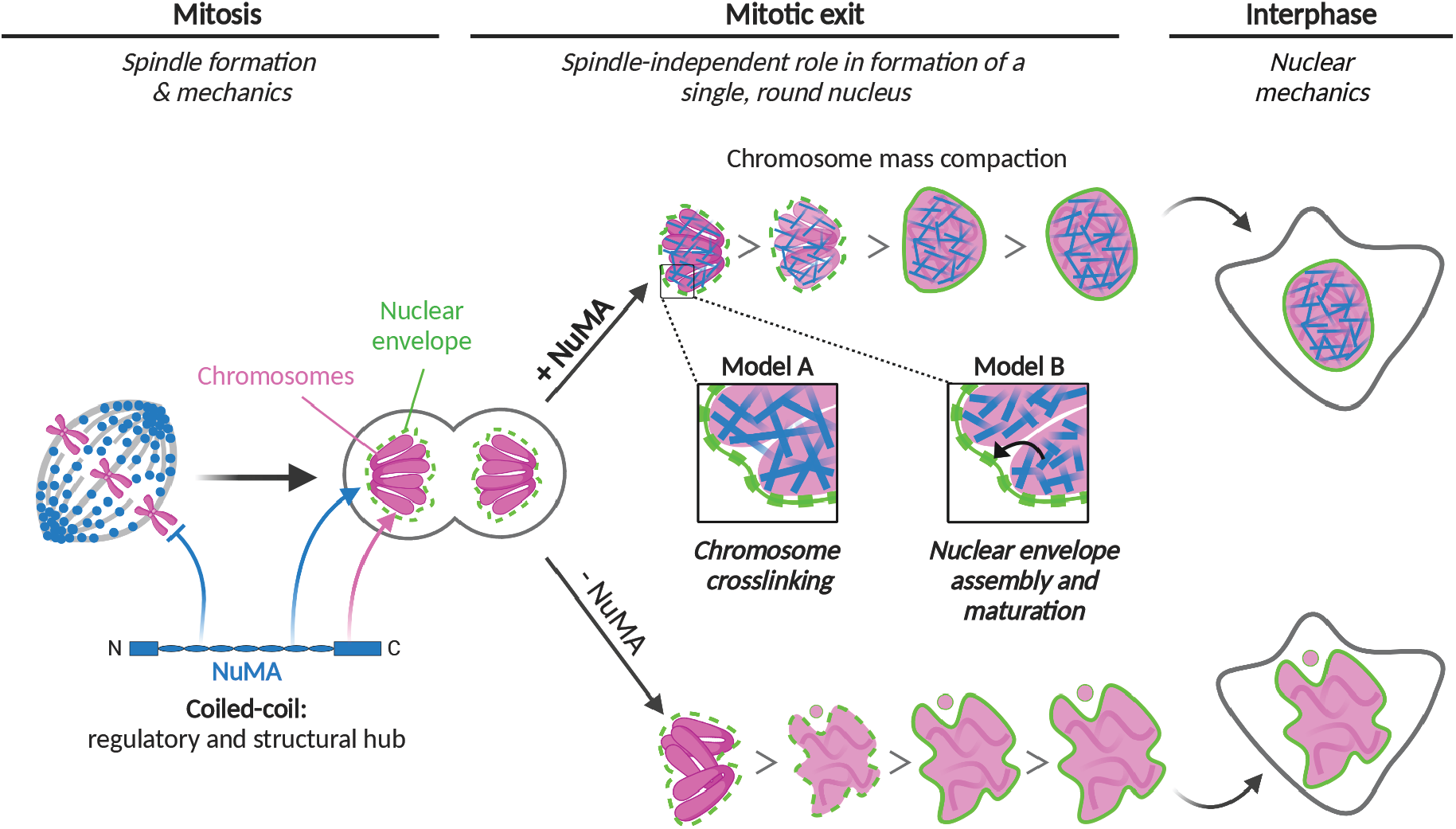
Validation of the RPE1 inducible Cas9 NuMA-KO cell line and of the nocodazole/reversine treatment. **(A)** Western blot (top) and immunofluorescence measurement (bottom) of endogenous NuMA levels in RPE1 cells after DOX-induced NuMA depletion for the indicated times. n=329 (0h), 448 (24h), 389 (48h), 327 (72h), and 223 (96h) cells. **(B)** Representative immunofluorescence images showing spindle morphology (tubulin), NuMA and chromosomes (Hoechst) in the indicated conditions. Cells were untreated or treated with nocodazole (664 nM) and reversine (320 nM) for 18-24h. Scale bar: 5 μm. Control cells assemble a spindle, NuMA KO cells assemble a perturbed spindle, and nocodazole and reversine-treated cells do not assemble a spindle. **(C)** Nuclear morphology in immunofluorescence images of DOX-induced Cas9 RPE1 cells not expressing sgRNAs. n=209 (no DOX control) and 328 (DOX) cells. **(D)** Representative immunofluorescence images of nuclear shapes in control and NuMA KO cells after DOX-induced NuMA depletion for the indicated times. The green line outlines the nucleus. Scale bar: 10 μm. **(E)** Nuclear solidity of interphase cells with respect to endogenous NuMA levels after DOX-induced NuMA depletion for 0-96 h. n=61 (1.2-1.8), 183 (0.8-1.2), 191 (0.4-0.8) and 214 (0.0-0.4) cells. Mann-Whitney test: ***p=0.0009; *p=0.01; ns, non-significant.

**Figure S2.**
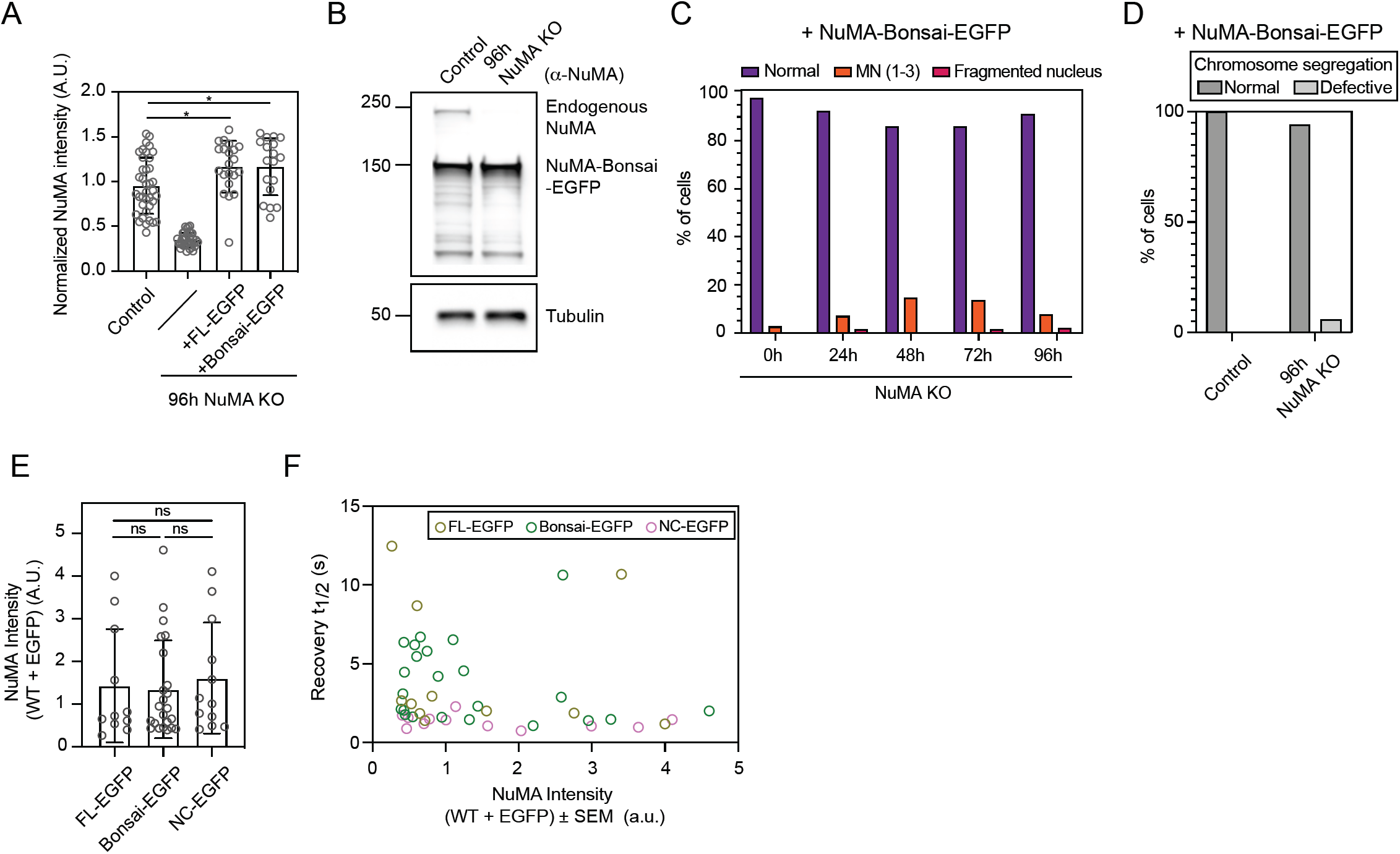
Validation of the RPE1 NuMA KO line stably expressing NuMA-Bonsai-EGFP and estimate of NuMA-EGFP intensity in cells used in the FRAP analysis. **(A)** Normalized NuMA intensity in uninduced NuMA KO RPE1 cells (control), NuMA KO cells and NuMA KO cells overexpressing the rescue constructs NuMA-FL-EGFP and NuMA-Bonsai EGFP. An antibody against the N-terminus was used in order to detect both endogenous and overexpressed protein. **(B)** Western blot of endogenous NuMA and NuMA-Bonsai-EGFP in uninduced NuMA KO RPE1 cells (control) and 96h NuMA KO cells stably overexpressing NuMA-Bonsai-EGFP. **(C)** Percentage of cells with different nuclear phenotypes at 0h, 24h, 48h, 72h and 96h of endogenous NuMA KO in cells stably overexpressing NuMA-Bonsai-EGFP and fixed for immunofluorescence as in Fig. 1B. n=205 (0h), 177 (24h), 181 (48h), 181 (72h) and 177 (96h) cells. **(D)** Chromosome segregation defects in live control and NuMA KO cells stably overexpressing Bonsai-NuMA-EGFP. Normal segregation indicates anaphase progression without lagging chromosomes or anaphase bridges. n=8 (control) and 17 (NuMA KO) cells. One out of 17 NuMA KO cells exited mitosis with a defective nucleus, and without completing anaphase, and was scored as segregation defective. **(E)** NuMA-FL-EGFP, NuMA-Bonsai-EGFP and NuMA-NC-EGFP fluorescence intensity in the cells analyzed in the FRAP experiment presented in Fig. 3D-E. n=11 (NuMA-FL-EGFP), 23 (NuMA-Bonsai-EGFP), and 12 (NuMA-NC-EGFP) Mann-Whitney test: ns, non-significant. **(F)** Fast recovery halftime as a function of NuMA-FL-EGFP, NuMA-Bonsai-EGFP and NuMA-NC-EGFP intensity in the cells from Fig. 3D-E. n=11 (NuMA-FL-EGFP), 23 (NuMA-Bonsai-EGFP), and 12 (NuMA-NC-EGFP).

**Figure S3.**
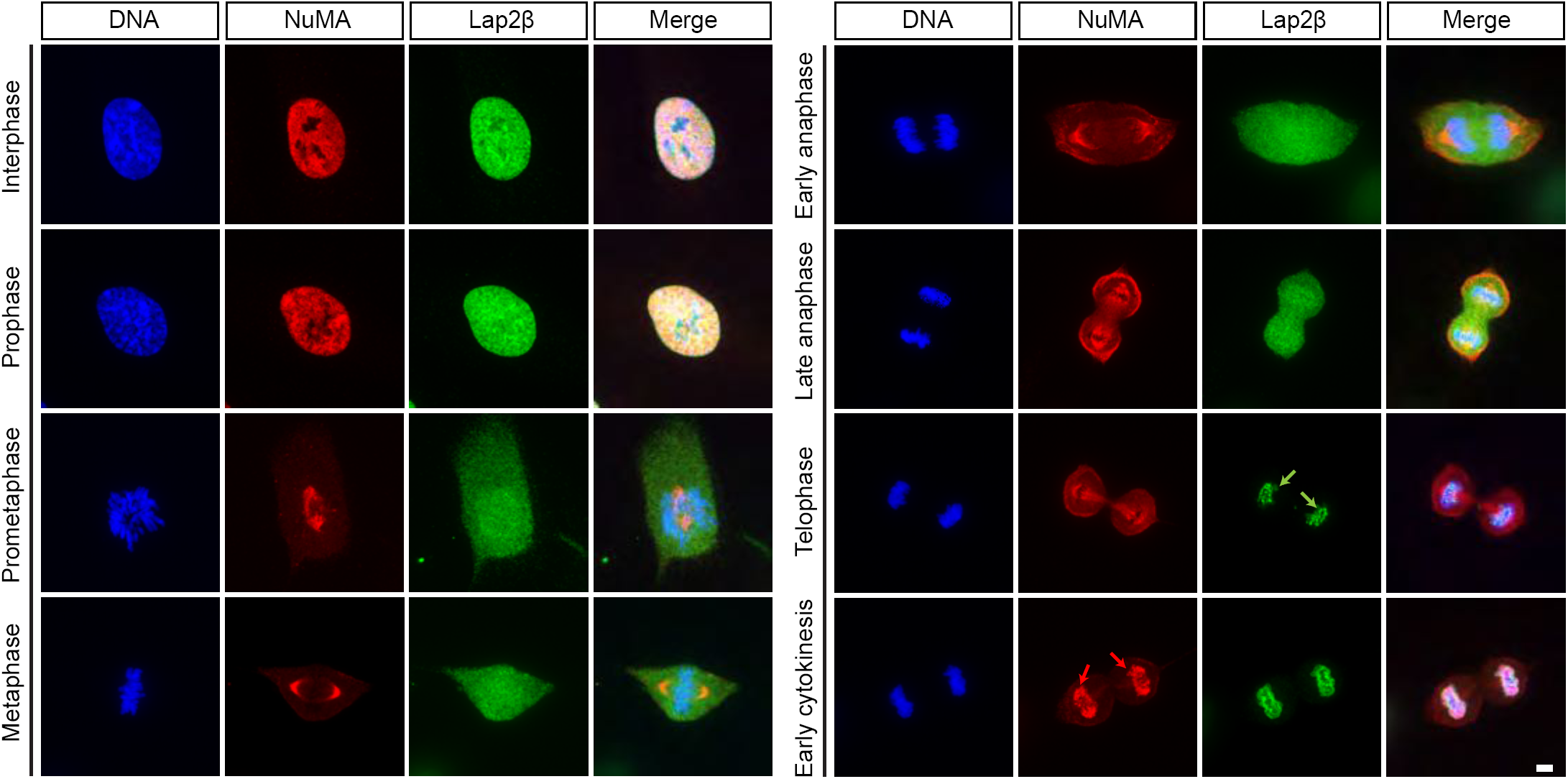
NuMA is only detectable on chromosomes after initial Lap2β recruitment. Representative immunofluorescence images of uninduced RPE1 NuMA KO cells showing localization of the nuclear envelope protein Lap2β (green), NuMA (red) and DNA (Hoechst, blue) over the cell cycle. Lap2β is detectable around the chromosome mass (green arrows) before NuMA appears on chromosomes (red arrows). Scale bar: 5 μm.

**Figure S4.**
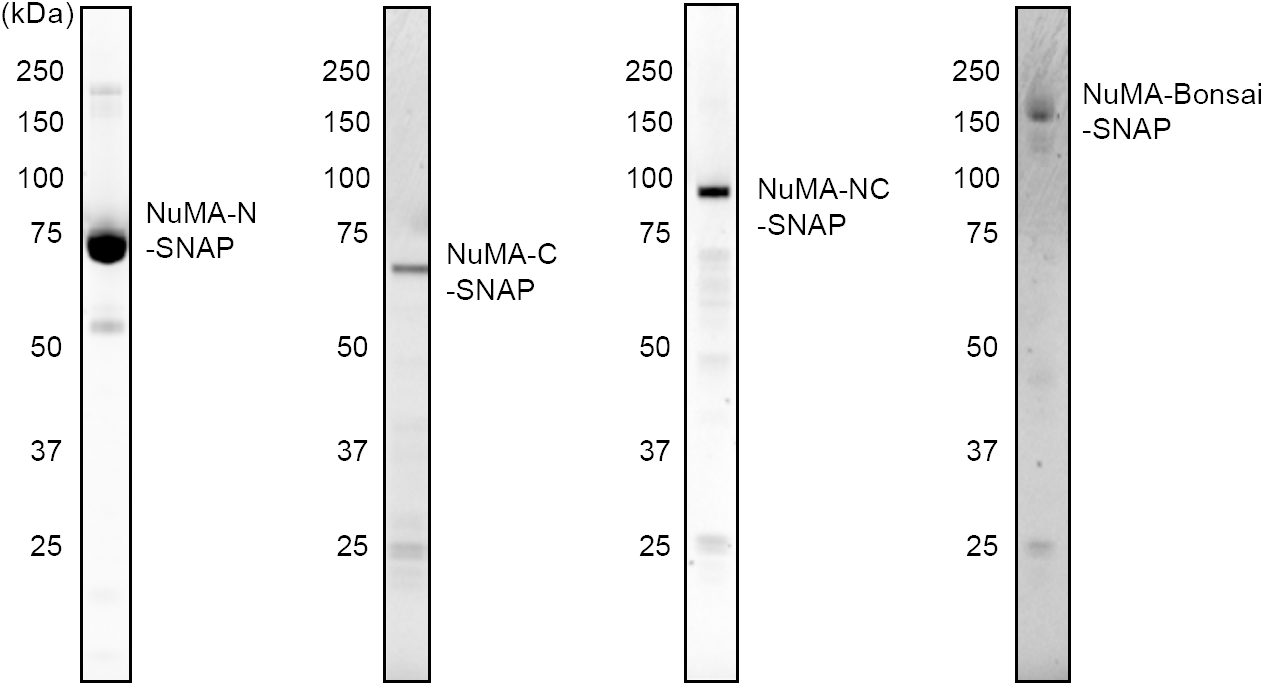
Purified proteins used for Electrophoretic Mobility Shift Assays. Coomassie blue stained gels with SNAP-tagged NuMA-N, NuMA-C, NuMA-Bonsai and NuMA-NC proteins purified from insect Sf9 cells.

**Figure S5.**
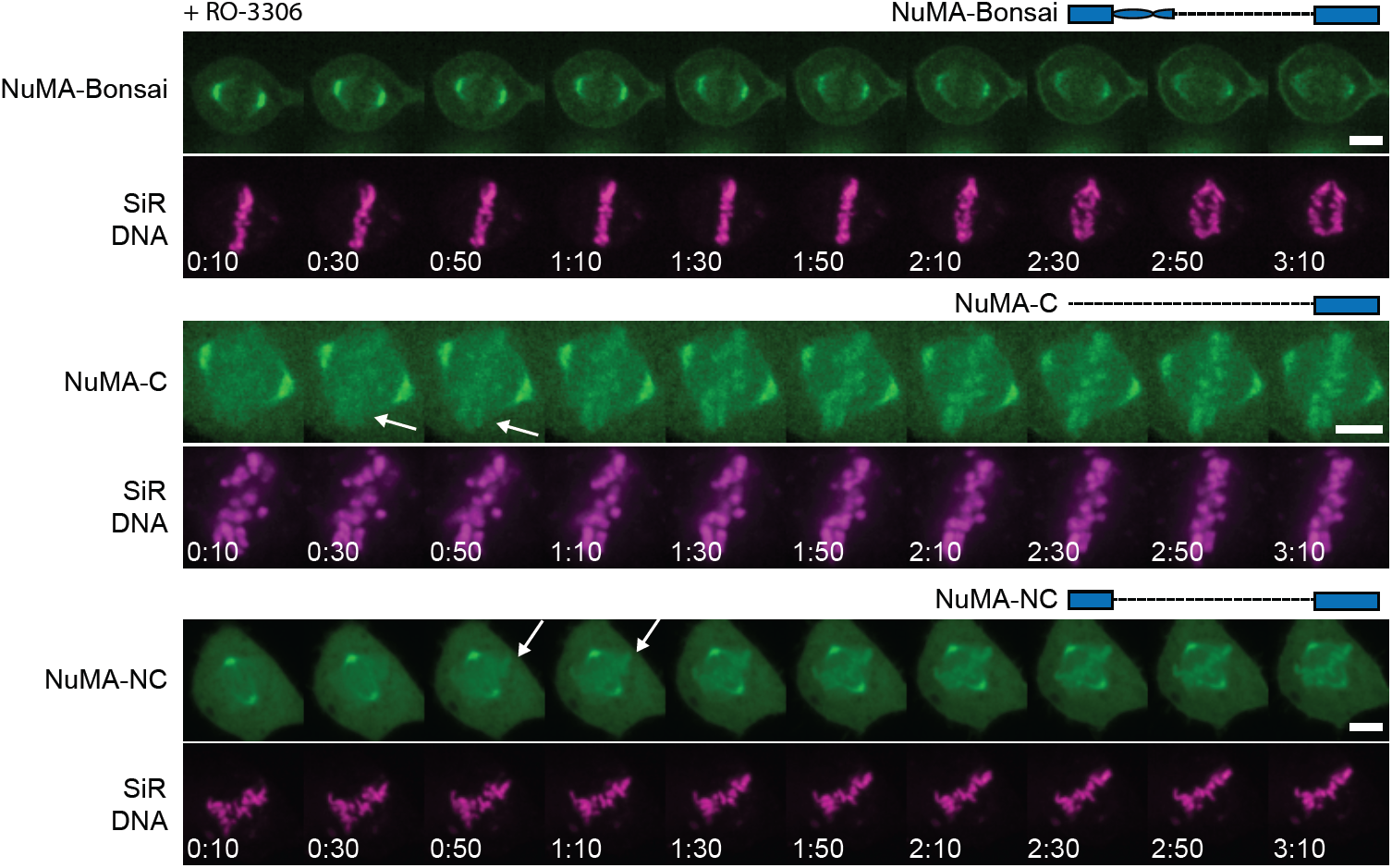
CDK1 activity does not affect NuMA binding to chromosomes during mitosis. Representative timelapse images of uninduced NuMA KO RPE1 cells stably expressing NuMA-Bonsai-EGFP, EGFP-Zdk1-NuMA-C or NuMA-NC-EGFP, treated with SiR-Hoechst (DNA), and treated with the CDK1 inhibitor RO-3366 approximately 10 s before 0:00. Time in min:s. Cells were imaged during metaphase to monitor the localization of the different EGFP-tagged NuMA truncation proteins on chromosomes. Arrows indicate the time at which we detect the tested EGFP-tagged NuMA truncations on chromosomes upon CDK1 inhibition. Scale bars: 5 μm.

## Video legends

**Video 1. Nuclear defects arise at nuclear envelope formation as NuMA-depleted cells exit mitosis.** Timelapse spinning disk confocal imaging of representative control (Video 1A) and NuMA-KO (Video 1B) RPE1 cells stably expressing mCherry-H2B (left, magenta) EGFP-Lap2β (center, green) and entering and exiting a spindle-less mitosis (nocodazole/reversine). Cells enter mitosis with a single and round nucleus, and exit with nuclear defects, which appear during nuclear envelope reformation. Time in min:s, with the frame corresponding to nuclear envelope breakdown set to 00:00. The start of nuclear envelope reformation becomes detectable at 00:12 in Video 1A and 0:24 in Video 1B. Scale bar: 10 μm. Videos correspond to the still images in Fig. 2D.

**Video 2. Overexpressed NuMA-FL-EGFP can form stable cable-like structures in the nucleus.** Timelapse imaging of a representative RPE1 cell transiently overexpressing NuMA-FL-EGFP imaged before and during fluorescence recovery after photobleaching (FRAP). The red circle highlights the photobleached area. Fluorescence recovery in the bleached area is minimal, consistent with NuMA-FL being able to form higher order structures in the nucleus. Time in min:s, with the frame corresponding to bleaching set to 00:00. Scale bar: 5 μm. Video corresponds to the still images in Fig. 3G.

**Video 3. NuMA-C has a higher affinity than NuMA-N for chromosomes in the nucleus.** Timelapse spinning disk confocal imaging of a representative RPE1 cell stably expressing EGFP-Zdk1-NuMA-C (left) and NuMA-N(1-705)-mCherry-LOV2 (right), before (<0:15), during (0:15-0:44, “+ Light”) and after (>0:45) exposure to blue light (488 nm laser). Upon blue light illumination, NuMA-C remains associated with chromosomes, while NuMA-N becomes diffusive. This indicates the NuMA-C binds to chromosomes in the nucleus with higher affinity than NuMA-N. Time in min:s. Scale bar: 5 μm. Video corresponds to the still images in Fig. 4B.

**Video 4. NuMA’s coiled-coil prevents its C-terminus from binding chromosomes at mitosis.** Timelapse spinning disk confocal imaging of a representative non-induced NuMA KO RPE1 cell transiently expressing NuMA-FL-EGFP, or stably expressing NuMA-Bonsai-EGFP, EGFP-Zdk1-NuMA-C or NuMA-NC-EGFP (in this order, green, left) and labelled with SiR-Hoechst (magenta, center) transitioning from metaphase to anaphase. NuMA-FL-EGFP and NuMA-Bonsai-EGFP, which contains part of the coiled-coil, do not bind chromosomes at mitosis, while NuMA-C and NuMA-NC, which lacks the entire coiled-coil, do. These indicate that NuMA’s coiled-coil regulates NuMA-C’s binding to chromosomes in mitosis. Time in min:s, with the frame corresponding to anaphase onset set to 00:00. Scale bar: 10 μm. Video corresponds to the still images in Fig. 4E.

